# A Toxic Tau-PFKFB3 Circuit Reduces F2,6BP Levels and Drives Neurodegeneration

**DOI:** 10.64898/2026.05.28.728580

**Authors:** Santi M. Mandal, Anirban Chakraborty, Shandy Shahabi, Mikita Mankevich, Adrian Povo-Retana, Tapan Biswas, Sergio Sanchez-Garcia, Sravan G. Sreenivasmurthy, Lucia Zhou Yang, Joseph Herdy, Jerome Mertens, Fred H. Gage, Johannes CM Schlachetzki, Balaji Krishnan, Lisardo Bosca, Tapas Hazra, Gourisankar Ghosh

**Affiliations:** Department of Biochemistry and Molecular Biophysics, University of California San Diego, La Jolla, CA 92093, USA; Department of Internal Medicine, University of Texas Medical Branch, Galveston, TX, 77555, USA; Instituto de Investigaciones Biomédicas Alberto Sols-Morreale (CSIC-UAM), Arturo Duperier 4, 28029 Madrid, Spain, and Centro de Investigación Biomédica en Red de Enfermedades Cardiovasculares (CIBERCV), Institute of Health Carlos III (ISCIII), 28029 Madrid, Spain; Department of Neurology, University of Texas Medical Branch, Galveston, TX, 77555, USA; Department of Neuroscience, University of California San Diego School of Medicine, La Jolla, CA 92093, USA; The Salk Institute, La Jolla, CA 92093, USA

**Keywords:** F2, 6-bisphosphate (F2, 6BP), Neurodegeneration, PFKFB3, PP2CA, Tau aggregation, DNA strand breaks

## Abstract

Alzheimer’s disease (AD) and related dementias are progressive neurodegenerative disorders manifested by aggregation of Tau and Amyloid beta (Aβ). Emerging evidence suggests that metabolic dysregulation contributes to AD pathogenesis, yet how metabolic alterations interface with neuronal integrity remains unclear. Here, we identify dysfunction in PFKFB3-F2,6BP (fructose-2,6-bisphosphate) metabolic axis as a key feature of AD. We show that pathological Tau aggregates aberrantly sequester PFKFB3, limiting its activity and resulting in F2,6BP depletion. F2,6BP exerts protective effects through multiple convergent mechanisms: (i) direct activation of polynucleotide kinase 3’-phosphatase (PNKP) to facilitate DNA strand break repair; (ii) transcriptional upregulation of the protein phosphatase 2A catalytic subunit (PP2CA) to limit Tau phosphorylation; (iii) stabilization of PFKFB3 to diminish its sequestration into aggregates; and (iv) direct inhibition of Tau aggregation. These findings establish F2,6BP as a central node linking metabolic regulation to both genomic stability and proteostasis in AD. Importantly, exogenous F2,6BP supplementation rescues multiple pathological features across diverse model systems, including induced neuronal cell lines (iN), primary neurons, organotypic hippocampal slice cultures, and in a *Drosophila* model of AD. These findings redefine F2,6BP as a metabolite that directly coordinates genome maintenance and proteostasis in neurons. Overall, this study identifies the PFKFB3-F2,6BP axis as a central driver of AD pathogenesis and a promising therapeutic target.

**Highlights:**

- Tau aggregates sequester PFKFB3 depletes neuronal F2,6BP
- F2,6BP links metabolism to DNA repair and Tau proteostasis
- F2,6BP activates PNKP and upregulates PP2A to counter Tau pathology
- F2,6BP supplementation rescues AD phenotypes across models

## Introduction

Alzheimer’s disease (AD) encompasses a spectrum of heritable and sporadic neurodegenerative disorders characterized by their progressive cognitive and motor decline (Langerscheidt et al., 2024) Jack et al., 2019). Hallmark pathological features include extracellular amyloid-β plaques (Aβ) plaques and intracellular Tau tangles. Aβ plaques arise from aberrantly processing of amyloid precursor protein (APP), generating aggregation prone 42-residue Aβ peptide (Chen et al., 2017). In parallel, Tau aggregation is driven by abnormal phosphorylation at multiple residues by kinases such as glycogen synthase kinase 3β (GSK3β), protein kinase A (PKA), and cyclin-dependent kinase 5 (CDK5) (Noble et al., 2003; Wegmann et al., 2021), and is further exacerbated by reduced activity of protein phosphatases (Manoharan et al., 2024; Zhou et al., 2009). In addition to these canonical features, AD is also marked by chronic neuroinflammation, mitochondrial dysfunction and widespread metabolic dysregulation (Heneka et al., 2025; Nadalutti et al., 2022; Riley and Tait, 2020; van Horssen et al., 2019; Wu et al., 2021; Wyss-Coray, 2016; Yin et al., 2016). However, how these diverse pathological features are mechanistically integrated remains poorly understood.

Metabolic dysfunction, particularly impaired glycolysis, is recognized as an early feature of AD (Zhang et al., 2021). A central regulator of glycolytic flux is fructose 2,6-bisphosphate, a potent allosteric activator of phosphofructokinase-1 (PFK1) and an inhibitor of fructose-1,6-bisphosphatase (Van Schaftingen and Hers, 1981; Van Schaftingen et al., 1981). F2,6BP is synthesized by the PFKFB (6-phosphofructo-2-kinase/fructose-2,6-bisphosphatase) family of bifunctional enzymes, among which PFKFB3 possesses a high kinase-to-phosphatase activity ratio, making it the principal enzyme for fructose-2,6-bisphosphate (F2,6BP) (Li et al., 2018; Macut et al., 2019). PFKFB3 is essential for organismal survival as *Pfkfb3*-deficient mice are embryonically lethal (Chesney et al., 2005). Importantly, PFKFB3 localizes to both the nucleus and mitochondria and associates with the DNA end-processing enzyme polynucleotide kinase 3’-phosphatase (PNKP) (Chakraborty et al., 2026a; Chakraborty et al., 2024).

PNKP removes 3’-phosphate (3’-P) termini from DNA strand break, an essential step in both single-strand and double-strand break repair pathways (Jilani et al., 1999; Karimi-Busheri et al., 1999; Shimada et al., 2015). Loss of PNKP activity in embryonic lethal and most embryos failing to survive beyond embryonic day nine In contrast, neural-specific deletion leads to postnatal mortality by day five, emphasizing its critical importance in both development and neurogenesis (Dumitrache and McKinnon, 2017; Shen et al., 2010). PNKP plays a direct and central role in neurogenesis by coordinating multiple DNA repair pathways to preserve genomic stability (Shimada et al. 2015).

Neurodegenerative disorders such as spinocerebellar ataxia type 3 (SCA3) and Huntington’s disease (HD), the 3’-phosphatase activity of PNKP is severely compromised (Chakraborty *et al*., 2024; Chakraborty et al., 2020; Gao et al., 2019; Poulton et al., 2013). We recently demonstrated that F2,6BP activates the 3’-phosphatase activity of PNKP in nuclear extracts prepared from Huntington’s disease and amyotrophic lateral sclerosis (ALS) patient tissues (Chakraborty et al., 2026b). These findings suggest that disruption of the PFKFB3-F2,6BP axis may represent a shared metabolic vulnerability across multiple neurodegenerative diseases. However, whether and how this pathway contributes to AD pathogenesis remains unknown.

Here, we identify dysfunction of the PFKFB3-F2,6BP axis as a central contributor of Alzheimer’s disease. We show that pathological Tau aggregates sequester PFKFB3, limiting its enzymatic activity and leading to depletion of F2,6BP. This metabolic deficiency drives a convergent pathogenic cascade, impairing DNA repair, promoting Tau hyperphosphorylation, and enhancing protein aggregation. Moreover, we demonstrate that F2,6BP directly modulates Tau aggregation and regulates multiple interconnected pathways governing proteostasis and genomic stability. Together, our findings uncover a previously unrecognized mechanism linking metabolic dysfunction to neurodegeneration and establish F2,6BP as a potential therapeutic target in Alzheimer’s disease.

## Results

### The 3’-phosphatase activity of PNKP is deficient in AD

To determine whether metabolic dysfunction impact AD, we quantified F2,6BP in hippocampal tissue extracts from 10 AD patients (Braak stage VI) and 10 controls (Braak I-II). F2,6BP levels were significantly decreased in AD (Braak VI) samples compared to controls (**Figure 1a & Supplementary Information**). To determine whether F2,6BP deficiency impairs DNA repair, we assessed the 3’-phosphatase activity of PNKP in nuclear extracts from cortical and hippocampal tissues. The assay measured the release of free phosphate from a radiolabeled 3’-phosphate-containing DNA strand within a gapped DNA duplex substrate (Chakraborty *et al*., 2024). Control reactions were supplemented with or without recombinant PNKP (rPNKP) and as expected, rPNKP catalyzed the release of free phosphate, whereas no product was detected in its absence. Nuclear extracts from AD cortex and hippocampus of a subset of samples displayed significantly lower 3’-DNA phosphatase activity of PNKP compared to controls (**Figures 1b and 1c)**. This result was confirmed in a larger number of cortical tissue samples (n=10 AD and n=10 control), with statistical analysis confirming significant differences (**Figure 1d).** Collectively, these findings indicate that reduced F2,6BP levels in the brains of AD patients contributes to the loss of PNKP activity, aligning with our finding that binding of F2,6BP to PNKP is necessary to maintain optimum 3’-phosphatase activity (Chakraborty *et al*., 2026a).

**Fig 1.**
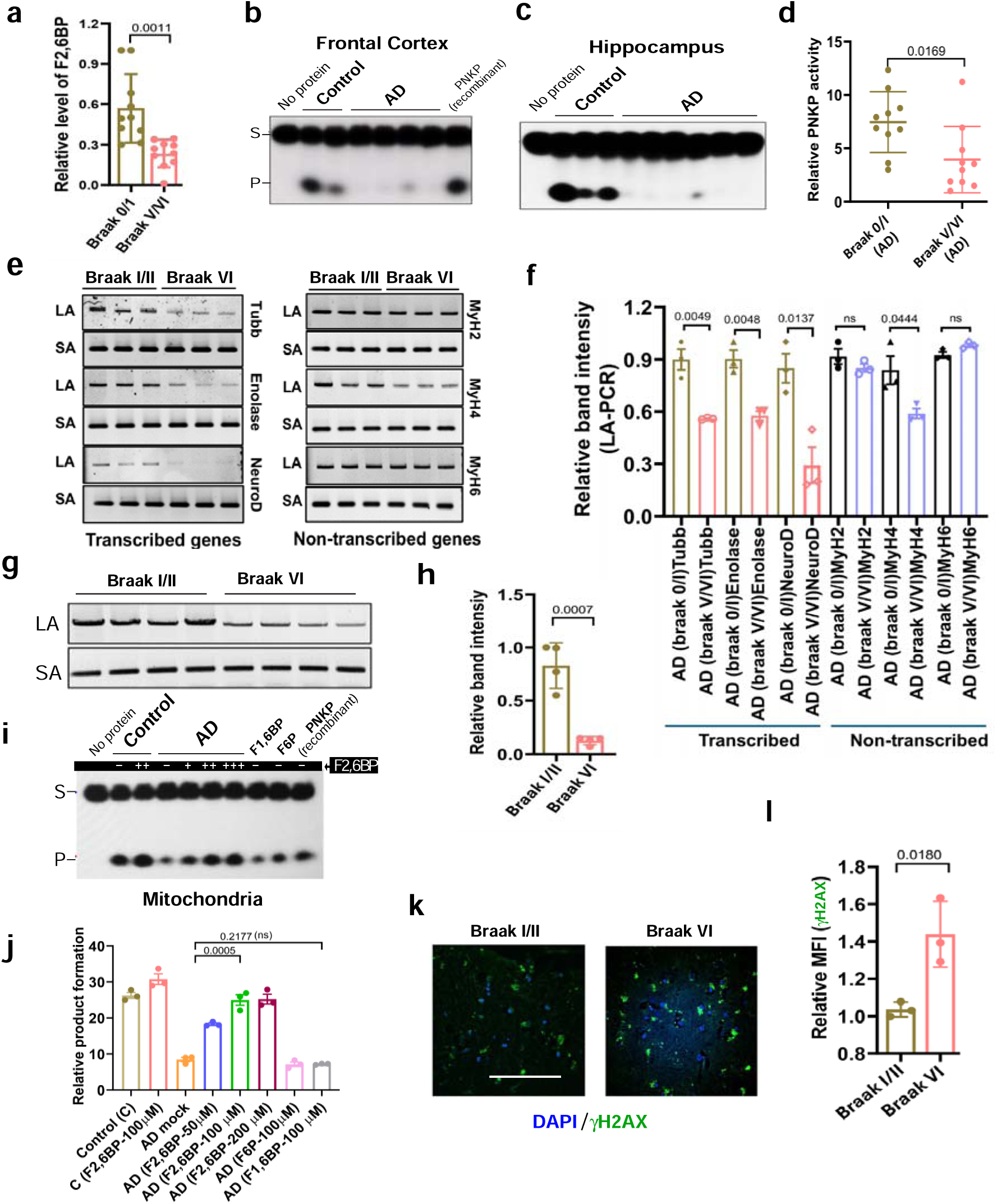
Levels of F2,6BP, loss of PNKP activity, elevation of DNA strand breaks and restoration of PNKP activity in AD and tauopathy patient samples. **a.** Bar graph showing relative levels of F2,6BP in human postmortem brain tissue across Braak stages I through VI (AD). Data are presented as mean ± SEM. Statistical significance (p value) was determined by two-tailed Student’s t-test comparing each among control and AD group. Sample size of Braak stage 0/I [n=10] and V/VI[n=10]. **b.** DNA 3’-phosphatase activity assay of PNKP in the nuclear extracts of frontal cortices from representative control and AD patient samples on a ^32^P-labeled 3’-phosphate-containing oligo substrates mimicking SSBs. 2 ng pure recombinant PNKP was used as a positive control. S= substrate and P= released phosphate. **c.** 3’-phosphate release activity of PNKP in the nuclear extracts of hippocampi of representative control and AD patient samples. **d.** Quantification PNKP activity from a larger cohort of AD patient and control samples (n=20-patients and 12-controls) expressed as relative activity. **e.** Representative agarose gel image showing long (LA) and short (SA) amplicon PCR products of nuclear DNA isolated from the frontal cortex of control and Alzheimer’s disease (AD) patients. The data indicate a significant elevated DNA repair deficiency in actively transcribing genes compared to non-transcribing regions. **f.** The normalized relative band intensities (LA/SA) are represented in the bar diagram with control samples arbitrarily set as 1 (n=4). Samples from AD, CBD and PSP patient and control samples were tested. Relative band intensity represents the mean ±SEM and p values are calculated following two-tailed Student’s t-test comparing each among control and AD group. **g & h.** Representative agarose gel image showing LA and SA amplicon PCR products (amplified using mitochondria-specific primers) of mitochondrial DNA isolated from the frontal cortex of control and AD patients. Mitochondrial DNA damage is significantly higher level of mitochondrial DNA damage in AD samples compared to the control group. Quantitation of band intensity (h). **i.** 3’-phosphatase activity of PNKP and its restoration by F2,6BP in mitochondrial extracts of AD patients and controls. NP (no protein) and rPNKP (recombinant PNKP) served as negative and positive control, respectively. F6P and F1,6BP showed no effect on the PNKP phosphatase activity. Statistical significance was determined using Student’s t-test, and p values are presented top of the bar graph with their corresponding group. **j.** Quantitation of PNKP activity in the nuclear extracts after the addition of F2,6BP (n=5 patients and n=5 controls). **k.** Representative immunofluorescence images of control and AD cortices stained with anti-γH2AX antibody. Nuclei are counterstained with DAPI. Images were captured using a confocal microscope at 60× magnification. Scale bar: 50 µm. **l.** Quantification of mean fluorescence intensity (MFI) corresponding to panel *k*. At least one slide each from three individuals of each group were imaged (n= 3). Data are presented as mean values +/-SEM.

Next, we quantified nuclear DNA damage directly in AD and control samples using long amplicon PCR (LA-PCR). In this assay, a relative decrease in the amplification of long (∼8–10 kb) versus short (∼250 bp) DNA fragments indicates increased DNA damage, as lesions block polymerase progression, and long amplicons provide a higher probability of encountering damage in the template (Chakraborty *et al*., 2024). We selected both actively transcribing genes in the brain, such as *Tubb3*, *Enolase2 and NeuroD,* as well as non-transcribing genes, such as *MyH2, MyH4, MyH6* **(Figure 1e**). Quantification of long amplicons showed a clear reduction in amplification of transcribing genes in AD samples compared to controls, whereas the difference was not significant in the amplification of non-transcribing genes (**Figure 1f**). These results provide clear evidence that damage accumulates preferentially in transcribed genes, which is consistent with an earlier report (Dileep et al., 2023). We also measured mitochondrial DNA damage in AD and control samples by LA PCR using specific sets of primer pairs. Similar to the nuclear genome, the mitochondrial genome of AD patients showed higher levels of damage, blocking amplification, compared to control samples (**Figures 1g and 1h**). Collectively, these findings indicate that reduced F2,6BP levels impair PNKP catalytic activity, leading to defective repair of DNA strand breaks and thereby linking metabolic deficiency to defective DNA strand break repair in AD.

### F2,6BP rescues defective PNKP activity in AD patient samples

To determine whether exogenous F2,6BP can restore PNKP activity in AD samples, we supplemented mitochondrial extracts from three cortical tissues and three age-matched control tissues with increasing concentrations of F2,6BP (25 µM, 50 µM and 100 µM). PNKP activity was restored in a dose-dependent manner across all AD samples. In contrast, structurally related metabolites, F1,6BP and F6P, did not elicit similar effects (**Figure 1i)**. Representative data from one AD and one control sample, along with quantification across all three samples are shown in **Figures 1i and 1j**. We next examined PNKP activity in nuclear extracts derived from both frontal cortex and hippocampal tissues. Male and female samples were analyzed separately (***Figures S1a-S1f***). Analysis of cortical and hippocampal nuclear extracts from both male and female samples revealed reduced PNKP activity in AD tissues which was robustly restored in a dose-dependent manner upon F2,6BP supplementation, with no sex-specific differences observed. Together, these results demonstrate that F2,6BP robustly restores PNKP activity in both mitochondrial and nuclear compartments in AD patient samples.

We next assessed the extent of unrepaired DNA strand breaks in cortical tissue samples from AD patients by visualizing and quantifying γH2AX foci using confocal microscopy imaging and a highly specific anti-γH2AX antibody. A significantly higher mean fluorescence intensity (MFI) was found in AD samples compared to controls (**Figure 1k)**. Quantitative analysis confirmed a statistically significant increase in γH2AX signal in AD tissues (**Figure 1l**), indicating accumulation of DNA double-strand breaks. Consistent with these findings, similar loss of genome integrity has been observed in other neurological disorders, including Huntington’s disease (HD), spinocerebellar ataxia type 3 (SCA3), and amyotrophic lateral sclerosis (ALS) (Chakraborty *et al*., 2026a; Chakraborty *et al*., 2026b; Chakraborty *et al*., 2024).

### PFKFB3 is transcriptionally upregulated but functionally inactivated in AD

To determine why F2,6BP levels are reduced in AD despite its role in DNA repair, we examined regulation of its biosynthetic enzyme, PFKFB3. Analysis of publicly available RNA-seq datasets revealed an increase in *PFKFB3 mRNA* expression in Braak stage VI patients compared to controls, whereas PNKP and other DNA repair genes remained unchanged (**Figure 2a**). We further found that mRNA levels of PFKFB3 increased with disease severity (**Figure 2b**). Elevated mRNA levels would typically predict increased protein abundance and activity. Because PFKFB3 catalyzes the synthesis of F2,6BP, we next assessed its enzymatic activity by measuring F2,6BP production in cortical extracts from AD patients. Unexpectedly, the catalytic activity of PFKFB3 was significantly reduced in AD cortex compared to control, indicating a lack of correlation between mRNA expression and functional enzymatic activity (**Figure 2c**). Immunofluorescence analysis of AD and control cortical brain sections revealed no significant difference in fluorescence intensity (**Figures 2d and 2e**). The discordance between RNA and protein levels prompted further examination of PFKFB3 localization. Careful inspection of images revealed punctate PFKFB3 structure in AD tissues, in addition to its presence as a diffuse soluble distribution, suggesting increased sequestration of PFKFB3 into aggregates. The punctate structures were present in all samples across all Braak stages, though their morphology differed: more diffuse in controls, smaller and more compact in intermediate stages, and larger with distinct sharp edges in late-stage disease (**Figure 2d**). To test if PFKFB3 co-aggregates with Tau, the same sections were co-stained with both MC1 (a monoclonal antibody that recognizes Tau aggregates) and anti-PFKFB3 antibodies. Confocal images and quantitation of fluorescence intensity confirmed their association in patient-derived samples and Tau aggregates increased with disease progression (**Figures 2d and 2e).**

**Fig 2.**
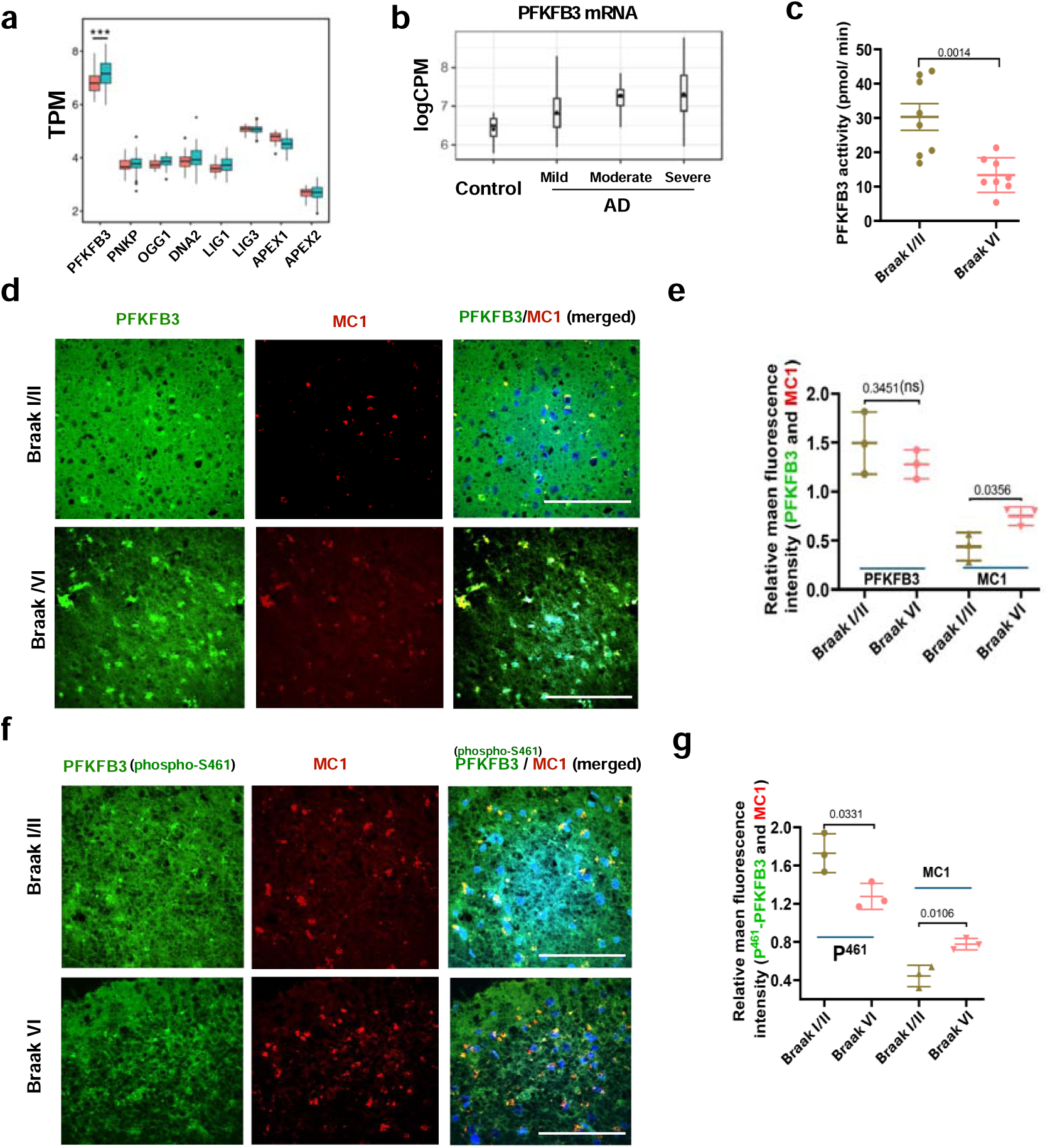
Co-aggregation of PFKFB3 and Tau in AD and tauopathy brains. **a.** Normalized gene expression levels, expressed as Transcripts Per Million (TPM) of several key DNA repair genes across two different Braak stages (Braak 0 and Braak VI) from a publicly available RNA-seq database. **b.** Normalized transcript levels of PFKFB3 at progressively severe disease stages (Braak stage 0= control; Braak stage I and II = mild; Braak stage III and IV = moderate; and Braak stage V and VI = severe) analysed from the same database as in panel A. **c.** Relative PFKFB3 enzyme activity in cortical samples from patients at Braak stages I/II and VI. Measurements were obtained from eight patient samples per group. **d.** Immunofluorescence images of cortical sections from AD samples stained with anti-PFKFB3 and MC1 antibodies at different Braak stages and compared (I/II vs VI). Images were acquired using a confocal microscope at 60× magnification. Three slides per group were analyzed, with multiple regions imaged on each slide. Scale bar: 50 µm. **e.** Quantitation of mean fluorescence intensity (MFI) of AD samples (I/II vs VI) for PFKFB3 and MC1. **f.** Immunofluorescence images of cortical sections from AD samples stained with anti–phospho-Ser461 PFKFB3 and MC1 antibodies at Braak stages I/II and VI. Images were taken on a confocal microscope at 60x magnification. Scale bar: 50 µm. **g.** Quantitation of mean fluorescence intensity of p-PFKFB3 and MC1 from different Braak stage of AD samples. Data are presented as individual data points with ± SEM.

Next, we examined the levels of PFKFB3 phosphorylation at Ser461, a modification known to enhance activity (Li *et al*., 2018) and its colocalization with MC1-positive aggregates. PFKFB3 Ser461 phosphorylation decreased with increasing severity and p-PFKFB3 was largely absent from MC1-positive puncta (**Figures 2f and 2g**). Consistent with this observation, Western blot analysis showed that phosphorylation of PFKFB3 at Ser461 was significantly reduced, whereas total PFKFB3 protein levels remained unchanged in patient versus control samples (**Figures S2a, S2b and S2c**). These results are in line with a report that showed in vitro-generated Tau fibrils interact with PFKFB3 in rat brain extracts (Ferrari et al., 2020).

Collectively, these results suggest that PFKFB3 is an essential factor maintaining genome integrity and metabolic homeostasis through F2,6BP synthesis. Loss of PFKFB3 phosphorylation at Ser461 correlates with its exclusion from Tau aggregates and reduced enzymatic activity, suggesting that phosphorylation may regulate its functional state and aggregation propensity. To compensate for reduced PFKFB3 enzymatic activity, neurons attempt to upregulate PFKFB3 transcriptionally. While translational regulation cannot be ruled out, our data support a model in which Tau-mediated sequestration of PFKFB3 limits its enzymatic activity despite increased transcription.

### F2,6BP prevents Tau aggregation in vitro

Given the central role of F2,6BP in regulating PNKP and PFKFB3, we asked whether it directly modulates Tau aggregation. We used recombinant full-length Tau and a truncated Tau expressed as a Myc fusion protein, Myc-TauK18 (Tau^K18^), that contains only the repeat domain (aa244-372) bearing the pathogenic P301S mutation (**Figure 3a**)(Iba et al., 2013; Jati et al., 2025). Tau^K18^ rapidly aggregates *in vitro* (within ∼2 hours), whereas full-length Tau (Tau^FL^) expressed as a poly-His fusion protein aggregates slowly (>12 hours) when incubated at 37 °C with gentle rotation (80 rpm). To test the effect of F2,6BP on Tau aggregation, Tau^K18^ was incubated with F2,6BP, F1,6BP or buffer at 37°C for 0 and 60 min followed by separation of soluble fraction from the total solution following centrifugation at 15,000 rpm for 20 min (***Figure S3a)***. Soluble fractions were analyzed by SDS-PAGE and Tau^K18^ was visualized by staining and quantified (***Figures S3b and S3c***). Insoluble pellets were assayed with Thioflavin T (ThT), which specifically binds β-sheet fibrils both in vivo and in vitro (Biancalana and Koide, 2010), to measure fluorescence (***Figures S3d & S3e***). In the presence of F2,6BP, aggregate formation was partially prevented, judged from enhanced soluble Tau^K18^ fraction and reciprocal decrease of insoluble Tau^K18^ in the pellet fraction in the presence of F2,6BP. F1,6BP showed a weaker effect (***Figures S3b-S3e)***. We next examined recombinant Tau^FL^ for aggregation. Tau^FL^ was incubated at 37°C with for 16 hr in the presence or absence of F2,6BP and continuous monitoring of ThT fluorescence in a TECAN plate reader. In the absence of F2,6BP, aggregates appeared after ∼12 hr, indicated by a sigmoidal increase in ThT fluorescence. In contrast, Tau^FL^ incubated with F2,6BP showed no fluorescence enhancement, indicating inhibition of aggregate formation (**Figure 3b**). This striking ability of F2,6BP to prevent Tau aggregation prompted us to test whether it could also disassemble preformed aggregates. Tau^K18^ was fluorescently labeled (FITC-conjugated), incubated at 37°C for 60 min to form aggregates, and then treated with F2,6BP, F1,6BP, or buffer while monitoring aggregation dynamics by live imaging for the next 60 min. While aggregates continued to grow in the absence of ligand, F2,6BP slowed aggregation and in some instances promoted partial disaggregation (***Figure S3f***).

**Fig 3.**
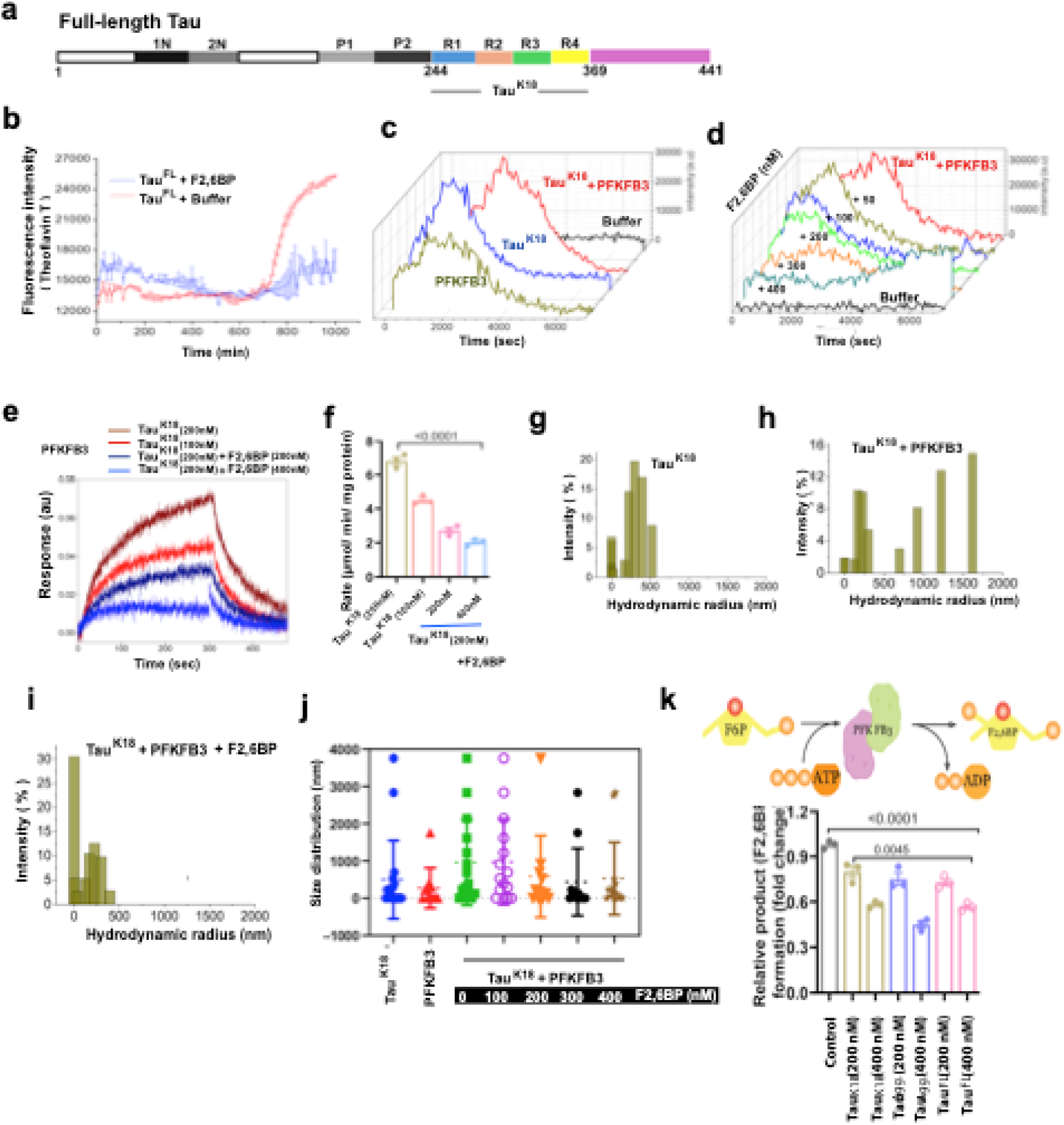
PFKFB3 associates with Tau-K18 and full-length Tau (F-Tau) in aggregates and reverse by F2,6BP. a. Schematic of full length 2N4R Tau (Tau^FL^) with repeat domains (R1-R4) and C-terminal end shown in colors. The Tau^K18^ fragment spans residues 241 to 372. b. Aggregation kinetics of F-Tau monitored by Thioflavin T (ThT) fluorescence. Tau aggregates slowly, and F2,6BP prevents aggregation. Mean ThT fluorescence intensity was recorded over 1000 minutes. The increase in fluorescence intensity reflects the rate of Tau aggregation, which is significantly reduced in the presence of F2,6BP. Data represent the mean ± SEM of three independent experiments. c. PFKFB3 aggregates independently and in associates with TauK^18^ aggregates. The time vs ThT fluorescence intensity plot showing co-aggregation of Tau^K18^ and PFKFB3 forming a distinct delayed aggregated species. d. Decrease in ThT fluorescence intensity with increasing concentrations of F2,6BP, implyes reduced formation of Tau^K18^:PFKFB3, PFKFB3 or Tau^K18^ aggregates. e. Binding kinetics between immobilized His-PFKFB3 and two concentrations of Tau^K18^ aggregates (100 and 200 nM) in the absence or presence of F2,6BP in a BLI instrument. f. Association rates between His-PFKFB3 and Tau^K18^ in the absence or presence of 200 and 400 nM F2,6BP. g-j. Effect of F2,6BP on the K18 aggregate formation measured by dynamic light scattering (DLS). g, h & i. Plots of hydrodynamic radius (Rh) vs % Intensity obtained from DLS measurement of Tau^K18^ (g), the Tau^K18^:PFKFB3 mixture (i), and the Tau^K18^:PFKFB3 mixture in the presence of 400 nM F2,6BP (i). Each sample was incubated for 60 min at 37°C before 20 consecutive DLS measurements were collected j. Distribution plot showing time-dependent changes of K18 aggregation as a function Rh under different conditions as described for g-i. k. K18, K18 (Agg) and full-length Tau inhibit F2,6BP synthesis by PFKFB3. The reaction schematic is shown on top. The reaction schematic (top) illustrates the assay design. Reactions were carried out for 12 hours at room temperature in the absence or presence of the indicated amounts of Tau proteins. F2,6BP levels were measured using the PPi-PFK1 assay as described in the Methods section. Bar graph shows relative product levels.

### F2,6BP prevents Tau-PFKFB3 aggregate formation by direct interaction

Given the potent ability of F2,6BP to prevent and reverse Tau aggregation, we next investigated whether it could also disrupt interactions between Tau aggregates and associated protein PFKFB3. Because PFKFB3 colocalizes with Tau inclusions in AD patient brains, we first examined whether Tau and PFKFB3 associate directly *in vitro*. Using the ThT fluorescence assay, we observed that both unaggregated and aggregated Tau^K18^, either alone or in complex with PFKFB3, produced strong fluorescence (**Figure 3c**), consistent with β-sheet–rich core structure formation. Interestingly, free PFKFB3 also showed ThT fluorescence, indicating an intrinsic capacity for self-aggregation. Incubation of Tau^K18^-PFKFB3 mixture with increasing concentrations of F2,6BP resulted in a dose-dependent reduction of ThT fluorescence. At 400 nM F2,6BP, ThT fluorescence was nearly abolished, indicating reduced formation of β-sheet-rich aggregates, consistent with inhibition of Tau and Tau-PFKFB3 co-aggregation (**Figure 3d**).

We further examined Tau-PFKFB3 interaction using a Biolayer Interferometry (BLI) assay. His-tagged PFKFB3 was immobilized on an anti-His biosensor, and Tau^K18^ was flowed over it to monitor association and dissociation. Aggregated K18 exhibited measurable binding to PFKFB3; however, this interaction was significantly diminished in the presence of F2,6BP, indicating that F2,6BP disrupts the association between the two proteins or prevents their association (**Figure 3e**). The apparent association rate decreased from 6.5 µmol/min to about 1 µmol/min in the presence of 400 nM F2,6BP (**Figure 3f**). These results support a model in which F2,6BP directly interacts with Tau, preventing its binding to accessory proteins such as PFKFB3 that are normally recruited into Tau inclusions.

To further substantiate these findings, we employed two additional complementary assays; dynamic light scattering (DLS) and gel electrophoresis. DLS measurements further confirmed the following findings: Tau protein aggregates displayed large hydrodynamic radius (∼300 to 400 nm) (**Figure 3g**), When Tau and PFKFB3 were mixed, a heterogeneous population ranging from 250 to 1600 nm was detected (**Figure 3h**). Addition of increasing concentrations of F2,6BP reduced the abundance of large species, and at 400 nm F2,6BP, the average size decreased to about 250 nm (**Figure 3i**). The comparative size distribution diagram further illustrated a concentration-dependent reduction in particle size with increasing F2,6BP levels (**Figure 3j**).

To visualize the PFKFB3-Tau coaggregation, we incubated PFKFB3 with preformed Tau^K18^ aggregates in the presence or absence of F2,6BP for 60 min and analyzed the soluble fraction by SDS-PAGE. Nearly all the PFKFB3 disappeared from the soluble pool with Tau^K18^ aggregates, suggesting coaggregation of PFKFB3 and Tau^K18^; however, F2,6BP treatment restored PFKFB3 solubility (***Figure S3g***). Similarly, when freshly prepared PFKFB3 and Tau^K18^ were incubated together, both proteins disappeared from the soluble pool after 60 minutes, but in the presence of 400 nM F2,6BP, remained soluble (**Figures S3h and S3i**).

To determine whether Tau association affects PFKFB3 catalytic activity, we measured F2,6BP synthesis in the presence of ATP and F6P. Addition of increasing concentrations of Tau^K18^, pre-aggregated Tau^K18^ or Tau^FL^, inhibited PFKFB3 activity in a dose-dependent manner (**Figure 3k**). These findings indicate that interactions between Tau aggregates and PFKFB3 are functionally harmful, leading to reduced enzymatic activity.

Previous studies have shown that Tau and other intrinsically disordered proteins interact through π–π stacking of arginine side chains (Ferrari *et al*., 2020). Our results suggest that F2,6BP interferes with these interactions, thereby preventing Tau–Tau and Tau∼Tau–PFKFB3 co-aggregation. Together, these findings demonstrate that a naturally occurring metabolite can directly modulate multiprotein aggregation driven by Tau, offering new mechanistic insight into Tau-mediated proteostasis dysregulation in neurodegeneration and suggesting potential therapeutic strategies.

### PP2A Suppression correlates with AD pathology

The observed Tau-PFKFB3 association prompted us to investigate the intrinsic role of PFKFB3 in the molecular mechanisms driving Tau aggregation. Tau aggregation is known to be induced by its hyperphosphorylation. Consistent with this, immunoblot analyses confirmed elevated phospho-Tau (p-Tau) in AD brains, whereas total Tau levels remained unchanged (***Figures S4a-Sd***). To investigate the mechanism underlying hyperphosphorylated Tau, we analyzed publicly available RNA-seq datasets from AD samples across different Braak stages (**Figure 4a**). Among several differentially expressed genes, we observed a progressive decline in expression of the catalytic subunit (PP2CA) and several regulatory subunits of protein phosphatase 2A (PP2A) with increasing disease severity, from Braak stage 0 to stage VI. Immunofluorescence analysis of cortical tissue sections confirmed reduced PP2CA (catalytic subunit) levels in Braak V/VII samples compared to Braak I/II controls (**Figures 4b and 4c**). Western blot analysis further validated these findings (**Figures 4d and 4e**). These results indicate that loss of PP2CA contributes to Tau hyperphosphorylation consistent with phosphatase-mediated dephosphorylation of Tau.

**Fig 4.**
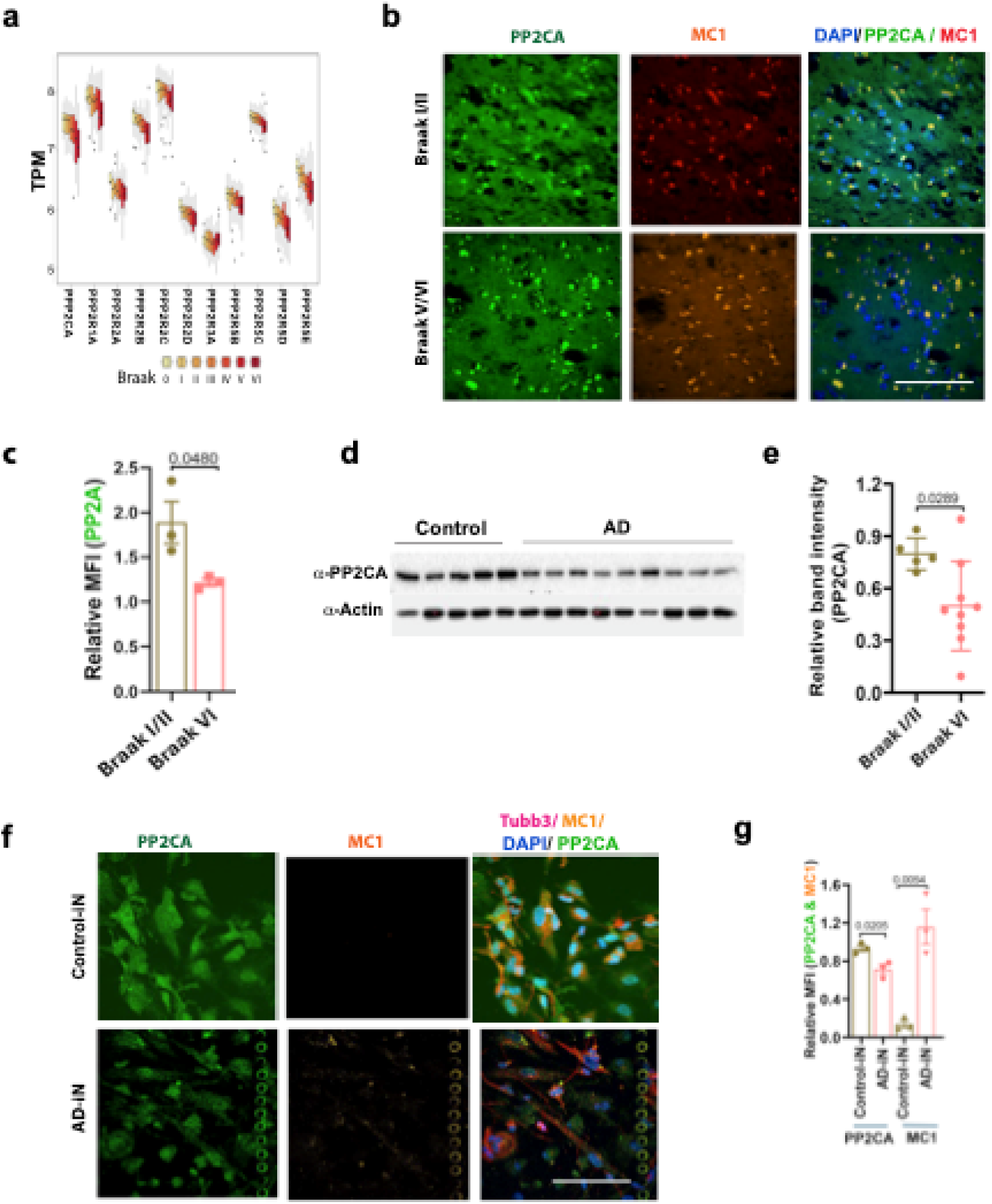
Decreased protein phosphatase 2A (PP2A) levels in AD, tauopathy patients, and AD patient-derived induced neurons (iNs) correlate with increased tau aggregation. a. Quantification of mRNA levels of the catalytic and regulatory subunits of PP2A at different Braak stages in AD cortical tissue samples. Data were obtained from publicly available RNA-seq database. b. Immunofluorescence images taken on a confocal microscope showing fluorescence intensity of cortical tissue sections from of Braak I/II and Braak VI patients samples stained with anti-PP2CA antibody. Scale bar: 50 µm. MC1 is used for Tau-aggregation specific antibody. c. Quantitation of mean fluorescence intensity (MFI) showing the difference of Tau aggregation between diseased and control samples (b). d. Western blot analysis was performed on cortical tissue extracts from control (Braak I) and AD (Braak VI) samples to assess the expression levels of PP2AC. e. Quantitation of band intensity in western blot h of both control and AD samples. f. Confocal microscopic images showing fluorescence intensity of AD- and control-iNs stained with three different antibodies as MC1 (tau aggregates), Tubb3 (neuron specific β-tubulin), and PP2CA (catalytic subunit alpha of protein phosphatase 2A), DAPI stained nuclei. Images were taken at 60x magnification. Scale bar: 50 µm j. Quantitation of relative PP2CA and MC1 levels based on their mean fluorescent intensity in control-iN and AD-iNs cells.

To further assess the relationship between protein PP2A and Tau aggregation, we examined PP2CA levels and Tau aggregation in induced neurons (iNs). iNs are differentiated neurons, generated from fibroblasts of AD patients, that recapitulate age-dependent pathology observed in sporadic AD, along with controls (Mertens et al., 2016; Mertens et al., 2015). Immunofluorescence imaging revealed a significant reduction of PP2CA levels in AD-iNs compared to control-iNs accompanied by increased MC1 staining, indicating enhanced Tau aggregation in the context of reduced PP2CA levels (**Figures 4f and 4g**).

### F2,6BP partially reverses toxic signatures in AD-iNs

The pronounced effect of F2,6BP in preventing Tau, PFKFB3 and Tau-PFKFB3 aggregation *in vitro* prompted us to test whether F2,6BP could also mitigate AD-associated toxicity. To this end, we assessed i) DNA damage, ii) PFKFB3 levels, iii) PP2CA levels, and iv) Tau aggregates in untreated and F2,6BP-treated AD-iNs by confocal microscopy. We tested three different iNs derived from different patients. Untreated AD-iNs exhibited elevated DNA damage compared to control-iNs as indicated by increased nuclear γH2AX mean fluorescence intensity (MFI) as assessed by confocal microscopy. Treatment of AD-iNs with F2,6BP markedly reduced the γH2AX fluorescence signal, restoring it to near-control levels (**Figures 5a and 5b**), consistent with reduced DNA damage.

**Figure 5:**
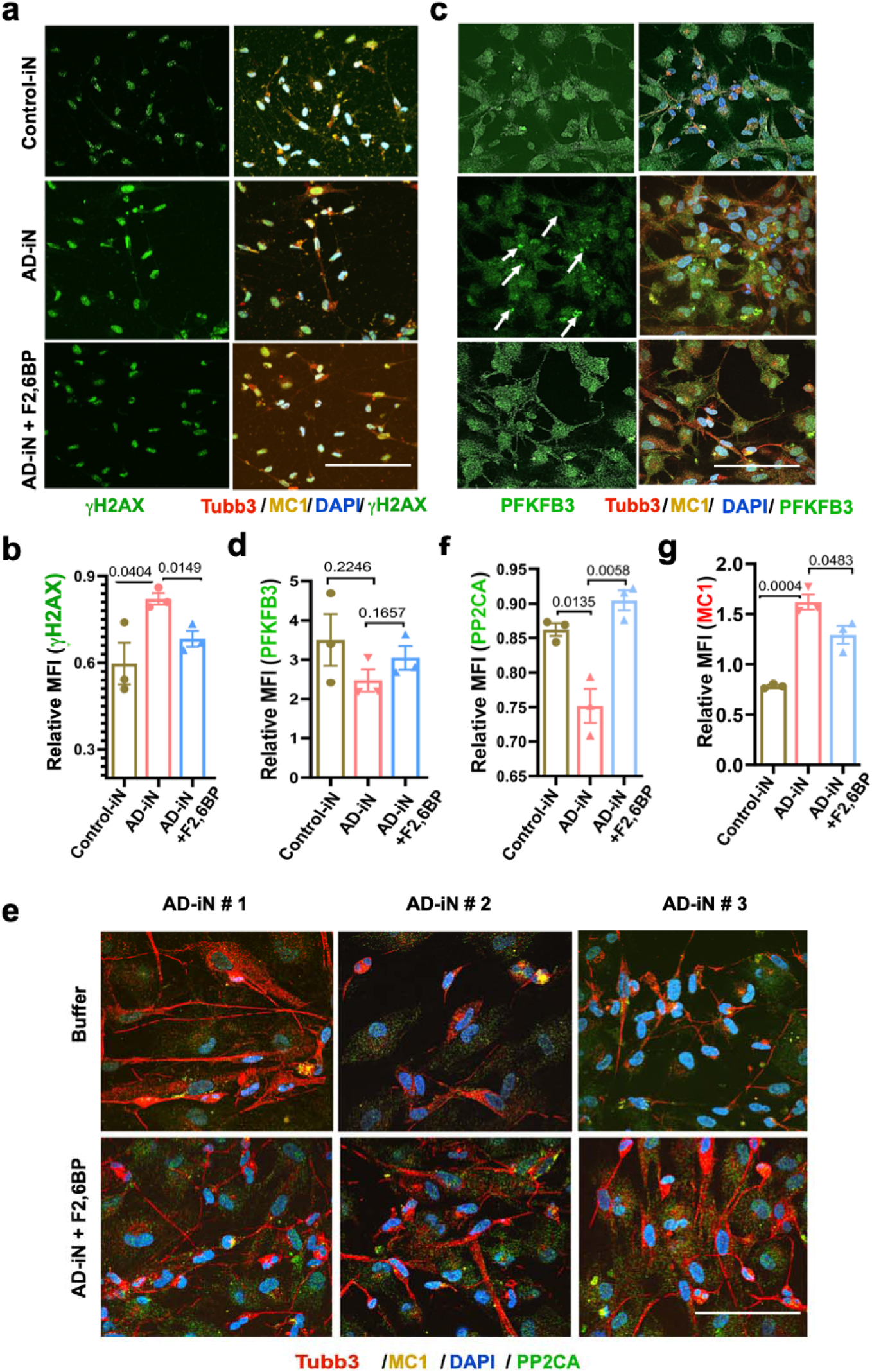
F2,6BP treatment reduces Tau pathology in AD-iNs by disrupting PFKFB3–Tau aggregates and increasing PP2A levels. a. Immunostained confocal images showing mean fluorescence intensity (MFI) of Con-iNs, AD-iNs and AD-iNs-treated with F2,6BP stained with 11-H2Ax, MC1, DAPI and Tubb3. Scale bar: 50 µm b. Quantitation of relative MFI of cells stained with 11H2Ax antibody demonstrating more DNA damage observed in AD-iNs and its recovery in the presence of F2,6BP. c. Similar experiments as in ‘a’ except cells stained with anti-PFKFB3 antibody. Arrows mark aggregates containing PFKFB3. d. Quantitation of relative MFI of experiments in c showing PFKFB3 levels under different growth conditions. e. Immunofluorescence of three different AD-iNs treated with or without F2,6BP stained with anti-PP2CA antibody. f. Quantitation of relative MFI showing PP2CA levels with and without F2,6BP treatment. g. Quantitation of relative MFI showing MC1 levels with and without F2,6BP treatment.

We next evaluated PFKFB3 protein levels in these cells and found PFKFB3 total protein levels were unchanged across AD-iNs, Con-iNs and AD-iNs, regardless of F2,6BP treatment (**Figures 5c and 5d**). However, AD-iNs exhibited prominent punctate PFKFB3 structures that were absent in con-iNs and reduced in treated AD-iNs suggesting F2,6BP prevents PFKFB3 aggregation. To test whether PFKFB3 puncta colocalize with Tau aggregates, we performed MC1 immunostaining. Tau aggregates were detected which colocalized with PFKFB3 puncta in AD-iNs (**Figures 5c and S5a**). In parallel, levels of Tau aggregates were also diminished (**Figures 5c and 5g).** We further assessed whether F2,6BP alters PFKFB3 transcription and found no enhancement of PFKFB3 transcript levels in AD-iNs treated with F2,6BP compared to untreated or treated with F1,6BP (**Figure S5b**).

We further investigated whether F2,6BP treatment modulates PP2CA levels. F2,6BP treatment consistently increased PP2CA levels across all three AD-iNs, as indicated by elevated MFI in confocal images (**Figures 5e and 5f**). Analysis of transcript levels revealed that while PFKFB3 mRNA levels remained unchanged, PP2CA mRNA levels were significantly increased in F2,6BP-treated cells, indicating transcriptional upregulation of PP2CA by F2,6BP, while F1,6BP had no effect (***Figures S5c and S5d***). These results indicate that F2,6BP deficiency contributes to reduced PP2CA transcription, PFKFB3 sequestration into Tau aggregates, diminished PNKP activity, and accumulation of DNA double strand breaks. Importantly, key pathological features observed in patient tissues, including defective DNA repair, reduced PP2CA protein levels, and Tau pathology, were recapitulated in AD-iNs and partially reversed upon F2,6BP treatment.

### Effect of exogenous F2,6BP on mouse hippocampal AD brain slices and *Drosophila* AD models

We postulated that supplementation of F2,6BP, given its diverse cellular functions, would provide greater neuroprotection in more complex models. To test this possibility, we used an organotypic hippocampal slice culture model in which human Tau aggregation is induced by seeding with exogenous Tau aggregates. In this system, hippocampal slices are transduced with AAV2-expressing human Tau carrying the P301S mutation (hTauP301S) and treated with 50 μM F2,6BP during Tau seeding or treated with buffer (**Figure 6a**).

**Fig 6:**
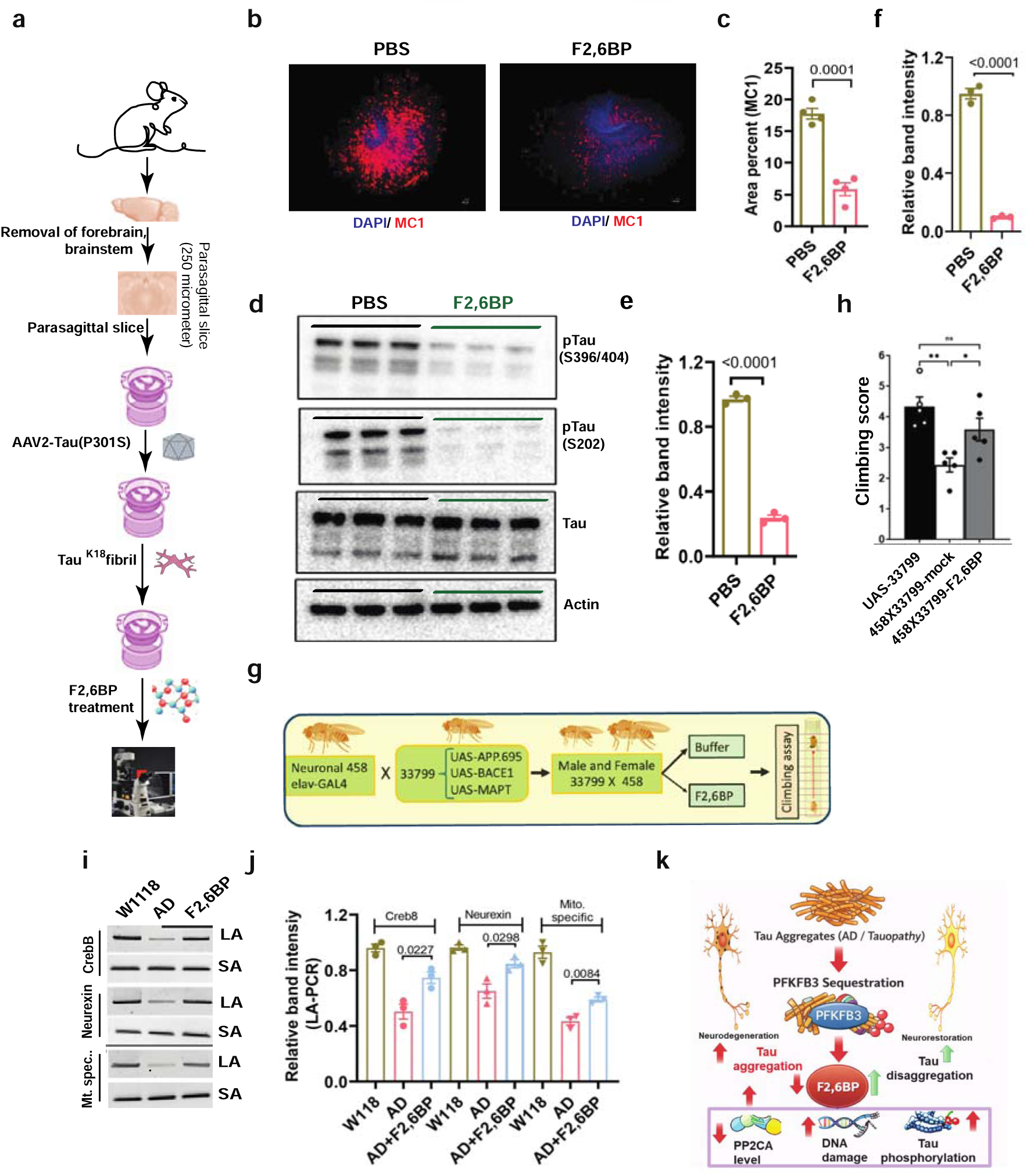
Ex>ogenous F2,6BP reduces tau aggregates in tauopathy slice culture and reverses pathology in Drosophila AD models. a. Schematic showing generation, treatment and imaging of organotypic hippocampal slice culture. b. Representative immunofluorescence images of hippocampal slices untreated (PBS) and treated with F2,6BP treated samples. Slices were stained with MC1(tau aggregates) and DAPI stained nuclei, showing significant reduction of tau aggregates in treatment. Scale bar: 1 mm. c. Quantification of relative percentage of MC1-stained area in PBS and F2,6BP-treated slices. Data were obtained from 4 different slices of individual set of experiments and presented as ± SEM d. WB analysis of total slice extracts using indicated antibodies of phosphor-Tau (S396/ 404) and S202. e&f. Quantification of specific phospho-Tau of S396/ 404 and S202 (f) western blot from three independent experiments. g. Schematics of AD fly generation and climbing assay. h. Quantitation of the climbing assay score. i. PCR analysis of the genomic DNA encompassing two transcribing genes and the mitochondrial genome. LA PCR amplifies 10 kb, and SA PCR amplifies 100 bp. j. Quantification of LA-PCR products of genomic DNA isolated from control fly, AD fly-untreated and AD fly-treated with F2,6BP k. A schematic drawing showing regulation of Tau aggregation–mediated sequestration of PFKFB3 drives metabolic dysfunction and neurodegeneration. Pathological Tau aggregates, characteristic of Alzheimer’s disease (AD) and tauopathies, aberrantly sequester the glycolytic regulator PFKFB3, leading to its functional inactivation. This results in depletion of fructose-2,6-bisphosphate (F2,6BP), a critical activator of glycolysis. Reduced F2,6BP levels contribute to multiple downstream pathological events, including decreased PP2CA levels, increased DNA damage, and enhanced Tau phosphorylation. These alterations collectively exacerbate Tau aggregation, forming a vicious cycle that promotes neurodegeneration. Conversely, restoration of PFKFB3 activity and F2,6BP levels may facilitate Tau disaggregation and support neuronal health restoration.

We then assessed Tau aggregation through detection of MC1-positive foci. Extensive aggregation was observed in buffer-treated control slices for two weeks following viral infection (**Figure 6b**). However, the extent of Tau aggregation was markedly reduced in F2,6BP-treated slices. Quantification of MC1-positive aggregates revealed nearly a 90% reduction in F2,6BP-treated hippocampal slices compared to untreated controls (**Figure 6c**). We also assessed Tau aggregation by monitoring phosphorylation at residues S202/204/205 and found a dramatic reduction of phosphorylation at these sites (**Figure 6d and 5f**). We found a similar protective role of F2,6BP in primary neurons infected with pathogenic Tau P301S mutant and seeded with Tau^K18^ oligomer. In F2,6BP-treated neurons, minimal MC1 staining was observed compared to buffer-treated control cells (**Figure S6**).

To further test if F2,6BP can counteract Tau pathology in a model organism, we utilized a 3X-TG AD model of *Drosophila* expressing APP.695, BACE1 and MAPT (BDSC# 33799) under pan-neuronal driver Elav Gal4. The AD flies exhibited impaired motor function (assessed by the negative geotaxis climbing assay). The experimental plan has been outlined in **Figure 6g**. Feeding these flies with a liquid diet with 50 μM F2,6BP for 21 days, significantly improved motor function in these flies compared to a diet with control buffer as measured by climbing scores (**Figure 6h**). We isolated DNA from control group and F2,6BP-treated Drosophila, tested DNA damage accumulation in nuclear DNA (nDNA) and Mitochondrial DNA (mtDNA) using gene-specific oligos by LA-qPCR. We indeed observed significant restoration of both nuclear and mitochondrial genome integrity in flies, supplemented with F2,6BP in the diet, which is consistent with their improved motor phenotype (**Figures 6i and 6j**). Therefore, these results suggest that repair deficiency in nuclear and mitochondrial genomes, consistent with dysfunction of the PFKFB3-F2,6BP-PNKP axis and exogenous F2,6BP can reverse such deficiency.

Collectively, F2,6BP acts through a network of convergent mechanisms to coordinate metabolic fidelity with neuronal resilience. First, it functions as a direct activator of PNKP, thereby promoting efficient repair of DNA strand breaks, a critical safeguard against genomic instability. Concurrently, F2,6BP drives transcriptional upregulation of PP2CA, enhancing phosphatase activity to restrain pathogenic Tau hyperphosphorylation. In parallel, it exerts functional stabilization of PFKFB3, effectively preventing its aberrant sequestration into Tau aggregates. Finally, F2,6BP acts directly upon the disordered Tau protein itself, inhibiting its aggregation in a manner independent of enzymatic regulation. Through this integrated, multi-modal axis, F2,6BP emerges not merely as a metabolic intermediate but as a master rheostat linking glycolysis to genome integrity, proteostasis, and AD suppression.

## Discussion

Our study identifies PFKFB3-F2,6BP axis dysfunction as a central event in the pathological cascade of Alzheimer’s disease (AD). We show that PFKFB3 is functionally impaired in AD brains. This impairment arises from its sequestration into Tau aggregates and reduced activating phosphorylation at Ser461, leading to decreased production of its metabolic product, F2,6BP. Depletion of F2,6BP triggers multiple interconnected effects: reducing PNKP activity, leading to impaired DNA repair; diminishing PP2A levels, resulting in Tau hyperphosphorylation; and directly promotes Tau aggregation, which enhances Tau–PFKFB3 sequestration. Together, these events form a self-perpetuating vicious cycle that ultimately drives neurodegeneration.

### A cascade of dysfunction triggered by PFKFB3 loss

Depletion of F2,6BP initiates a multifaceted cascade in neuronal dysfunction. Reduced F2,6BP compromises DNA strand break repair by impairing the activity of PNKP, which requires F2,6BP for optimal function (Chakraborty et al., 2026). DNA repair proteins are enriched at well-defined hotspots that protect essential genes. These hotspots are enriched with histone H2A isoforms and RNA binding proteins and are associated with evolutionarily conserved elements of the human genome (Reid et al., 2021). Additionally, diminished F2,6BP correlates with lower levels of protein phosphatase 2A (PP2A), a key regulator of Tau dephosphorylation. While F2,6BP also appears to influence transcriptional regulation, the mechanisms remain unknown. A key pathological driver is the sequestration of active PFKFB3 into Tau aggregates, resulting in a loss of its enzymatic function. The initiating mechanism remains uncertain, two interrelated possibilities may explain this: First, that PFKFB3 is intrinsically aggregation prone; second, oxidative modification promotes its interaction with Tau. It contains more cysteine residues in its unstructured C-terminal region which may undergo conformational changes upon oxidation increasing its affinity for Tau, promoting formation of unstable complexes that progressively aggregate.

### Structural basis of protein aggregation and F2,6BP-mediated protection

The ability of F2,6BP to both prevent and disassemble Tau aggregates is particularly intriguing. Since these effects were observed in vitro with purified full-length Tau^FL^ or the Tau^K18^ fragment, they likely result from direct interactions between Tau and F2,6BP, independent of other cellular components. Given that only F2,6BP, but not the closely related F1,6BP, exerts this effect, our data suggests a unique phosphate position-dependent conformational effects specific to F2,6BP.

We further demonstrate that PFKFB3 aggregates in vitro and colocalizes with Tau aggregates in patient samples. Our observation is consistent with a report that showed that PFKFB3 co-aggregates with Tau and other neuronal proteins *in vitro* (Ferrari *et al*., 2020). The driving forces for these associations are believed to involve π-stacking interactions mediated by arginine residues within intrinsically disordered regions (Ferrari *et al*., 2020). Although no detailed folding studies of PFKFB3 exist, unresolved regions in crystal structures (Cavalier et al., 2012) and the presence of six arginine residues in the C-terminal disordered domain support this model. Remarkably, F2,6BP can prevent and even reverse PFKFB3 aggregation and its association with Tau, suggesting a post-translational regulatory mechanism whereby F2,6BP stabilizes PFKFB3, maintaining its active pool and thereby sustaining its own synthesis, forming a positive feedback loop. Such reversible aggregation and inactivation are not unique to PFKFB3; several glycolytic enzymes undergo similar transitions during oxidative stress or nutrient deprivation (Bie et al., 2024; Itakura et al., 2015; Webb et al., 2017).

### PFKFB3 is a potential risk factor for AD

Reports on the role of PFKFB3 in neurodegeneration have been conflicting. Some studies suggest that PFKFB3 deficiency contributes to AD and other brain pathologies (Fu et al., 2015; Kwon et al., 2024; Wang et al., 2021), whereas others report the opposing effects (Burmistrova et al., 2019; Rodriguez-Rodriguez et al., 2012; Wang et al., 2023). Neuronal PFKFB3 is typically maintained at low levels through continuous degradation and its overexpression can induce neuronal death via excess reactive oxygen species (ROS), generated by hyperactive glycolysis (Almeida et al., 2010; Herrero-Mendez et al., 2009). Conversely, PFKFB3-driven glycolysis in neurons is important during brain excitation (Diaz-Garcia et al., 2017). We show that PFKFB3 is localized to mitochondria in neurons, consistent with our recent report that PFKFB3 supports mitochondrial health in striatal neuronal cells (Chakraborty *et al*., 2026a). Together, these observations suggest that an optimal threshold of PFKFB3 is required, with both deficiency and excess being detrimental. A recent study further demonstrated that aberrant PFKFB3 expression is a key driver of neurodegeneration in multiple sclerosis (MS) (Woo et al., 2025). Notably, MS is an autoimmune disease initiated by glial cells, unlike most other neurological disorders that originate primarily in neurons. How glia-initiated dysfunction triggers neuronal degeneration through PFKFB3 activation remains to be elucidated. Collectively, our observations and previous reports suggest that maintaining optimum PFKFB3-F2,6BP balance is needed for neuronal homeostasis, and that either excessive or insufficient amounts increase neuronal vulnerability. We, therefore, propose that PFKFB3 is a risk factor for AD.

### F2,6BP as a regulatory and therapeutic molecule

The bidirectional relationship between PFKFB3 and F2,6BP represents a previously unrecognized metabolic–structural feedback system: PFKFB3 generates F2,6BP, while F2,6BP, in turn, stabilizes PFKFB3 and prevents its aggregation. This mutual reinforcement loop positions F2,6BP as a critical regulator of neuronal homeostasis. Importantly, F2,6BP also demonstrates broad neuroprotective potential. In *Drosophila* models of multiple neurodegenerative diseases, spinocerebellar ataxia 3 (SCA3), Huntington’s disease (HD), amyotrophic lateral sclerosis (ALS), and AD, F2,6BP supplementation restored both physiological and behavioral functions. We previously demonstrated mitochondrial dysfunction, persistent DNA strand breaks, and impaired glucose metabolism constitute a shared pathological axis across neurodegenerative diseases. Here we extend this framework by identifying additional aberrations arising from F2,6BP loss mediated through Tau-PFKFB3 aggregation. Therefore, targeting F2,6BP-regulated pathways could offer a potentially unifying therapeutic approach. Small molecules such as epigallocatechin gallate (EGCG) have been shown to disaggregate Tau and other amyloid fibrils by direct binding (Andrich and Bieschke, 2015). EGCG binds to the amyloid scaffold of Tau fibril, producing conformational stress to break the paired helical structure (Andrich and Bieschke, 2015; Seidler et al., 2022). However, despite potent activities in vitro and in model systems, none of these fibril disassemblers have proven to be therapeutically effective. In contrast, F2,6BP supplementation reverses multiple pathological processes beyond simple Tau disassembly, suggesting that it acts on broader metabolic and signaling pathways to restore neuronal homeostasis.

A key unresolved question concerns the mechanism of F2,6BP cellular uptake. Decades of work with the related metabolite fructose-1,6-bisphosphate (F1,6BP) demonstrated that exogenous F1,6BP confers protection against ischemic injury in liver, heart, and lung by preventing oxidative lipid accumulation and sustaining ATP synthesis through oxidative phosphorylation (Antunes et al., 2006; Lazzarino et al., 1992; Sano et al., 1995; Xu et al., 2024). Radiolabeled tracer studies revealed rapid tissue uptake and metabolic conversion of F1,6BP within minutes, implying active intracellular transport rather than an exclusively extracellular mechanism (Hardin and Roberts, 1994). We made similar observations using fluorescently labeled F1,6BP and F2,6BP, demonstrating that both compounds enter cells. Although no specific transporter has yet been identified for either F1,6BP or F2,6BP, the evidence strongly suggests that such uptake pathways exist.

In conclusion, our findings identify dysregulation of the PFKFB3-F2,6BP catalytic cycle, driven by PFKFB3 dephosphorylation and co-aggregation with Tau, as a key pathogenic route underlying neuronal toxicity in AD. F2,6BP supplementation promotes Tau-PFKFB3 disaggregation, restores metabolic imbalance, and reverses multiple downstream defects. These findings highlight F2,6BP as a promising physiological modulator capable of restoring neuronal metabolism and mitigating Tau-associated toxicity in AD.

### Limitation of the Study

While our study establishes important roles for PFKFB3 dysfunction and F2,6BP depletion in AD, several important questions remain unresolved. First, we have not determined whether PFKFB3-F2,6BP dysregulation occurs at early stages of disease or a secondary, late-stage pathology. Second, although our data show reduced PP2A protein and RNA levels, it is not clear how F2,6BP regulate its transcription. Third, the observed discordance between PFKFB3 protein and RNA levels have not been mechanistically resolved. Fourth, the full spectrum of F2,6BP-binding targets is unknown, thus the broader biological consequences of altered F2,6BP availability remain to be defined. Fifth, although F2,6BP supplementation shows strong neuroprotective effect, it remains unclear whether chronic elevation of F2,6BP may produce detrimental outcomes in neurons, given known vulnerabilities to glycolytic overstimulation. Finally, we do not know how exogenous F2,6BP crosses plasma membrane. Future studies will be required to address these mechanistic uncertainties and to evaluate the translational feasibility and safety of F2,6BP-based therapeutic strategies.

## Materials and Methods

### Chemicals

All chemicals were purchased from Merck/Sigma/Aldrich, Roche, Invitrogen, and Bio-Rad.

### Antibodies

Commercial antibodies were from Invitrogen (phopho-PFKFB3 ref: PAS-114619; PFKFB3 ref: MA5-32766); Cell Signaling (Danvers, MA; vinculin ref: 13901; γH2Ax Ref: 6731; PP2CA ref: 2038); ThermoFisher (HT7-Tau Ref: NM1000,, phospho-Tau-AT8, ref: MN1020, and Strep-HRP, Ref: N100); Sigma (Tau ref: T9450; anti-rabbit-HRP Ref: A0545 and anti-mouse HRP Ref:A0168); Gifts from Albert Einstein College of Medicine-Peter Davies antibodies (MC1, PHF1 and CP13); Biobharati Lifescience, India (PNKP Ref: BB-AB0105 and Actin Ref: BB-AB0024) and Origene (Tubulin 3 Ref: TA425959)

### Preparation of human tissue extracts

Tissue samples were processed following standard operating procedures, with the appropriate approval of the Ethics and Scientific Committees. Frozen samples of brain cortex (20-40 mg) were homogenized at 4°C in 0.5 ml of 10 mM Tris-HCl (pH 7.5), 1 mM MgCl_2_, 1 mM EGTA, 1 mM DTT, 50 mM NaF, 0.5 mM phenylmethanesulphonyl fluoride, 10 μM leupeptin and a protease and phosphatase inhibitor cocktail (Merck). The extracts were centrifuged for 15 min at 13,000***g*** and the supernatants were aliquoted (50 μl) and stored at −20°C. Protein concentration was determined using the detergent-compatible Bio-Rad reagent.

### Preparation of F2,6BP from tissue samples

Brain tissue (20–40 mg) or > 5 × 106 cells were homogenized in an alkaline buffer and subjected to heat treatment at 80°C for 10 min to denature endogenous enzymes. The samples were then rapidly cooled and neutralized to pH 7.0–7.5 using 0.1 M HEPES/acetic acid mixture. Finally, the extracts were centrifuged at 13,000 × g for 10 min to remove insoluble debris. The neutralized tissue extract is added to the reaction mixture, and F2,6BP activity is determined by monitoring the decrease in absorbance at 340 nm.

### Synthesis of F2,6BP

A reaction cocktail was prepared consisting of 60 mM Tris-HCl (pH 7.5), 1.5 mM DTT, 5 mM Potassium Phosphate (pH 7.5), 20 mM KCl, 40 μM EDTA, 6 mM MgCl_2_, 5 mM ATP, 1 mM F6P, 10% Glycerol and1 mg/mL BSA in 200μL. The reaction was initiated by adding 100 μg PFKFB3 and incubated at 37°C for 90 mins followed by quenching the reaction with 50 μL 1M NaOH and heating at 80°C for 5 mins. The mixture was centrifuged to remove any precipitate followed by dilution to 2 mL with 10 mM TEABC (pH 8.5). The diluted reaction mixture was applied to MonoQ column pre-equilibrated with 10 mM TEABC (pH 8.5). F26BP was eluted using 20 to 35% gradient with 800 mM TEABC as buffer B. Peak fractions were pooled based on the phosphate release assay by PNKP, dried and dissolved in 20 mM Tris-Cl (pH=8.0). F2,6BP was also synthesized chemically in the Bosca laboratory (Bosca et al., 1985; Mojena et al., 1992), and the chemically synthesized compound was used in some early PNKP assays.

### Assay of 6-phosphofructo-2-kinase/fructose-2,6-bisphosphatase (PFK2/BPase-2) activity

Poly(ethylene)glycol 6000 was added to aliquots of tissue extracts to reach 5% concentration (weight/volume). After centrifugation at 7000***g*** for 15 min more poly(ethylene)glycol was added to the supernatant to reach 15% to fully precipitate the PFK2/FBPase-2 protein. After resuspension of the pellet in the extraction medium, PFK2/FBPase-2 activity was determined at 30°C and pH 8.5 with 5 mM MgATP, 5 mM Fru-6-P and 15 mM Glc-6-P. Samples were collected at different times (10 μl) and one volume of 50 mM NaOH at 80°C was added to stop the reaction. One unit of PFK2/FBPase-2 activity is the amount of enzyme that catalyzes the formation of one pmol of Fru-2,6-P_2_ per min (Bosca et al., 1988; Bosca *et al*., 1985).

### 3’-Phosphate release Assay

The 3’-phosphatase activity of PNKP in the nuclear extract of post-mortem patient frontal cortex/cerebellum and age-matched control subjects (2.5 µg) or with purified recombinant PNKP (2 ng) was conducted as we described previously^19^. Five pmol of the radiolabeled substrate was incubated at 37°C for 15 min in buffer A (25 mM Tris-HCl, pH 8.0, 100 mM NaCl, 5 mM MgCl_2_, 1 mM DTT, 10% glycerol and 0.1 μg/μl acetylated BSA). Nuclear extracts were prepared following the protocol used for Co-IP studies. For kinase activity assay, γP32 labeled ATP was incubated in kinase assay buffer (80 mM succinic acid pH 5.5, 10 mM MgCl_2_, 1 mM DTT, 2.5% glycerol) along with 1.0 μg/μl acetylated BSA, and 0.6 pmole labeled substrate for 30 min at 30°C^33^. 100 fmol of PNKP and 2.5 pmole of cold substrate were used in this assay. For *in vitro* PNKP restoration/abrogation, similar assays were done after incubation of F2,6BP/F6P/F1,6BP (in amounts as indicated in the main text and figure legends) with the nuclear extracts for 15 min. The radioactive bands were visualized in PhosphorImager (GE Healthcare) and quantitated using ImageQuant software. The data were represented as % product (phosphate) released from the radiolabeled substrate with a value arbitrarily set at 100%.

### Long amplicon quantitative PCR (LA-qPCR)

Genomic or mitochondrial DNA was extracted using the Genomic tip 20/G kit (Qiagen) per the manufacturer’s protocol, to ensure minimal DNA oxidation during the isolation steps. The DNA was quantitated by Pico Green (Molecular Probes) in a black-bottomed 96-well plate and gene-specific LA qPCR assays were performed as described earlier^18,19,41,42^ using Long Amp Taq DNA Polymerase (New England BioLabs). A different set of transcribed (neuronal differentiation factor 1 (NeuroD), tubulin β3 class III (TUBB) and gamma-enolase (Enolase) vs. non-transcribed (muscle-specific myosin heavy chain 2 [MyH2) and MyH4 genes were used for the LA-qPCR assay. The LA-qPCR reaction was set for all genes from the same stock of diluted genomic DNA sample, to avoid variations in PCR amplification during sample preparation. Preliminary optimization of the assays was performed to ensure the linearity of PCR amplification with respect to the number of cycles and DNA concentration (10-15 ng). The final PCR reaction conditions were optimized at 94°C for 30 s; (94°C for 30 s, 55-60°C for 30 s depending on the oligo annealing temperature, 65°C for 10 min) for 25 cycles; 65°C for 10 min. Since amplification of a small region is independent of DNA damage, a small DNA fragment (∼200-400 bp) from the corresponding gene(s) was also amplified for normalization of amplification of the large fragment. The amplified products were then visualized on gels and quantitated with ImageJ software (NIH). The extent of damage was calculated in terms of relative band intensity with a control siRNA/mock-treated sample considered as 100. All oligos used in this study will be listed in the Supplementary Table 1.

### His-Tau (full length) and Myc-K18 purification and in vitro aggregate formation

Full length wild type Tau as a poly-histidine tagged protein and the repeat domains (K18) of Tau bearing the P301L as a Myc fusion protein was expressed in E. coli BL21 (DE3) strain. Cells at OD A600 ∼0.6 were induced with IPTG (100 μM) at 30 °C for 3.5 h, pelleted and the pellet was resuspended in the lysis buffer (500 mM NaCl, 20 mM MES, pH 6.8, 1 mM EDTA, 0.2 mM MgCl2, 1 mM PMSF, 1 mM DTT), and sonicated for lysis. The supernatant was heated at 90 °C for 20 min. was clarified by centrifugation and was dialyzed overnight (Buffer A: 50 mM NaCl, 1 mM MgCl2, 0.1 mM PMSF, 2 mM DTT, 1 mM EGTA, 20 mM MES pH 6,8). Subsequently, cation exchange (HiTrap SP HP, 5 ml column from Cytiva) was performed to purify K18. Protein was eluted by a salt gradient. Pure proteins were concentrated and further dialyzed against assay buffer (PBS pH 7.4, 1 mM DTT and 2 mM MgCl2). Proteins were concentrated to 2 mg/mL and storred at −;80 C. Aggregates were formed by incubating the proteins at 37 °C with gentle shaking (2 hr for K18) and 24 hr for His-Tau^FL^ in the presence of 44μg/ml Heparin.

### Western blot analysis

Protein extracts from brain tissues were completed with 1% (w/v) Triton X-100. Proteins were resolved on SDS-PAGE gels and then transferred to nitrocellulose membranes. Protein bands were visualized using a Luminata chemiluminescence detection system (Merck Millipore) and an Image-Quant LAS 500 imager (GE Healthcare Life Sciences, Freiburg, Germany) and were quantified using ImageJ [National Institutes of Health (NIH), Bethesda, MD, USA]. The ratio between the intensities of each protein band and vinculin (used as reference) was calculated and plotted.

### Slice Culture

Organotypic hippocampal slice cultures were prepared from postnatal day 7-9 from WT pups as described previously (Croft & Noble, 2018). Briefly, pups were culled by decapitation and the hippocampus quickly dissected in oxygenated artificial cerebrospinal fluid (ACSF). 400 µm slices were cut using a tissue slicer. Approximately 18-24 slices were collected and cultured in Millicell culture inserts (Millipore) in 6 well plates (6-8 slices per insert). Hippocampal slices were transduced with AAV viral vector expressing P301S human Tau (2x10^11^/mL) for 24 hours. AAV was removed and K18 preformed Tau fibrils were added to the culture media (1.5 ug/mL) and incubated for one media change. Subsequent media changes were either media + PBS or media containing F2,6BP. Culture media was changed every 3 days supplemented with PBS or F2,6BP and the slices were harvested after 21 days in culture for Western blot or immunostaining.

### Primary neuron cultures

Primary cortical neuronal cultures were propagated using postnatal day 0 or 1 mice. The cerebral cortices of 3-4 pups were dissected on ice, meninges removed and placed in the cold dissection medium (DM) consisting of 6 mM MgCl2, 0.25 mM CaCl2,10 mM HEPEs (100X), 0.9% Glucose, 20 mM D-AP5 (Cayman, NC1368401), and 5 mM NBQX (Tocris Bioscience, 10-441-0). The tissues were digested with papain (Worthington, LK003176) in 37 C water bath for 20 min, followed by 5 min incubation with low OVO and DNase I incubation to stop the digestion. The digested samples were triturated and filtered through 70 mm cell strainer. The cell suspension was centrifuged at 1000rpm for 10 min, and the cell pellet was gently resuspended in 20 mL B27/NBM/ High glucose media and centrifuged again at 850 rpm for 5 min. The resulting cell pellet was resuspended in B27/NBM (1 mL/mouse brain). The cells were counted and plated onto the coverslips at 250,000 in 24-well plates for immunofluorescent imaging. Antibodies used: MAP2 (1:400, Millipore-Sigma Cat#AB-5622), MC1 (1:500, generated by Dr. Peter Davies and a received as a Gift from Albert Einstein College of Medicine).

### WT- and AD-iNs culture

‘iN-ready’ frozen fibroblasts regrown in in DMEM containing 15% tetracycline-free <u>fetal bovine serum</u> and 0.1% NEAA (Thermo Fisher) in the presence of puromycin (1 μg/mL, Sigma Aldrich). Cells were trypsinized and pooled into high densities (30.000 – 50.000 cells per cm^2^) and, after 24 h, the medium was changed to neuron conversion (NC) medium based on DMEM:F12/Neurobasal (1:1) for 3 weeks. NC medium contains the following supplements: N2 supplement, B27 supplement (both 1x; Thermo Fisher), doxycycline (2 μg/ml, Sigma Aldrich), <u>Laminin</u> (1 μg/ml, Thermo Fisher Scientific), dibutyryl-cyclic-AMP (500 μg/ml, Sigma Aldrich), human recombinant <u>Noggin</u> (150 ng/ml; Preprotech), LDN-193189 (5 μM; Fisher Scientific Co) and A83-1 (5 μM; Santa Cruz Biotechnology Inc.), <u>CHIR99021</u> (3 μM, LC Laboratories), <u>Forskolin</u> (5 μM, LC Laboratories) and SB-431542 (10 μM; <u>Cayman</u> Chemicals). For further maturation beyond 3 weeks, flow cytometry-isolated or -non-isolated iN cultures were switched to BrainPhys (STEMCELL Technologies)-based neural maturation media containing N2, B27, <u>GDNF</u>, <u>BDNF</u> (both 20 ng/ml, R&D), dibutyryl cyclicAMP (500 μg/ml, Sigma Aldrich), doxycycline (2 μg/ml, Sigma-Aldrich) and laminin (1 μg/ml, Thermo Fisher).

### Treatment of iN cells with exogenous metabolites

We treated iNs with 25 to 50 μM of F2,6BP or F1,6BP was mixed and incubated for few min to 48 hours for different experiments.

### Immunofluorescence

Paraffin-embedded brain slices pre-mounted on slides were de-paraffinized in xylene for 10 mins, followed by rehydration with sequential incubation in 100%, 90%, 70% EtOH, 5mins each. Sections were placed in Citrate buffer (pH 6) solution in a pressure cooker at low pressure for 10 minutes for antigen retrieval. The slides were blocked (2.5% Normal Goat Serum for 1 hr at room temperature) and incubated with primary antibody for overnight at 4 °C, followed by 3X PBS wash for 15 min each. Next, a secondary antibody and DAPI (1:2000, Thermo Fisher Cat #D3571) were added and incubated for an hour at room temperature.

### DLS analysis

Dynamic Light Scattering (DLS) analysis was performed using a DynaPro NanoStar (Wyatt Technology, USA) to monitor the time-dependent aggregation of K18 peptide fragments of the Tau protein. Samples were incubated with different supplements, including F1,6BP, F2,6BP, and the standard aggregation-promoting agent, heparin. The final concentration of K18 in the solution was 1 µM, while the concentrations of supplements ranged from 1 to 50 µM. Aliquots were collected at regular time intervals and analyzed. The hydrodynamic diameter and polydispersity index (PDI) were recorded using the instrument’s Dynamics software to assess changes in particle size distribution. A gradual increase in average particle size and scattering intensity indicated the progressive formation of K18 aggregates. Buffer (TBS, pH 7.5) and monomeric K18 samples were used as controls to ensure measurement accuracy. These results provided quantitative insight into the kinetics and stability of K18 aggregation over time.

### Human Transcriptome analysis

For human transcript analysis, normalized gene expression counts in the parahippocampal gyrus (Brodmann area 36) were obtained from Mount Sinai Brain Bank (MSBB) through the Synapse platform (syn16795937). A total of 9 individuals at BRAAK stage 0, 23 at BRAAK stage 1, 23 at BRAAK stage 5, and 64 at BRAAK stage 6 were included in the analysis.

### Continuous live imaging analysis of *In-Vitro* Tau aggregate

Fluorescence labeling of Tau protein was carried out using a maleimide-ester–based fluorescent compound designed to react with the amine groups of lysine residues. Purified Tau was incubated with the dye in TBS buffer (pH 7.5) at 37°C to enable efficient conjugation. Unreacted dye molecules were removed by dialysis using a 3 kDa cut-off membrane against fresh buffer to obtain the purified labeled protein. The degree of labeling and purity were verified by UV–Vis spectroscopy and SDS-PAGE. The resulting fluorescent Tau conjugates were subsequently used for continuous fluorescence imaging. Fluorescence-tagged K18 was aggregated for 2 h in TBS buffer (pH 7.5) at 37 °C, and aggregation was confirmed using a thioflavin T (ThT) assay. The resulting aggregates were then used as a substrate (1 µM) with or without F2,6BP to study the assembly and disassembly processes *in vitro* under controlled conditions. The reaction mixture was incubated in TBS buffer (pH 7.5) at 37 °C and monitored continuously for 10 hours using a fluorescence microscope equipped for time-lapse imaging. Changes in fluorescence intensity and distribution were recorded to observe the progressive association and dissociation of Tau aggregates.

### Biolayer Interferometry (BLI) analysis

Biolayer Interferometry (BLI) experiments were performed using an Octet system (ForteBio, USA) to investigate the interaction of full-length His-tagged Tau protein (0.2 µM). The Tau protein was immobilized onto Ni–NTA biosensors via its His-tag, ensuring stable attachment and proper orientation for binding analysis. Association and dissociation kinetics were measured upon exposure to fructose-1,6-bisphosphate (F1,6BP), fructose-2,6-bisphosphate (F2,6BP), and heparin at concentrations of 1 µM and 5 µM. Each analyte was allowed to interact with the Tau-coated sensor, followed by dissociation in buffer to record real-time binding profiles. The response curves revealed concentration-dependent differences in binding behavior, indicating distinct or opposing affinities of Tau toward the analyte molecules. All experiments were performed in TBS buffer (pH 7.5) at room temperature, and the data were analyzed using the Octet system software to determine kinetic parameters and binding strength.

### Drosophila maintenance and treatment

All Drosophila stocks were maintained at 25° C on standard fly food under a 12:12-h light-dark cycle. Drosophila strains were purchased from Bloomington Drosophila Stock Center (BDSC, Bloomington, Indiana, USA: UAS-HTT (#33808, RRID: BDSC 33808), expresses human Huntingtin (HTT) with long polyQ (glutamine) repeat of 128 amino acids, and Elav GAL4 (#458, RRID:BDSC_50892), shows pan neuronal GAL4 expression. BDSC 33808 were crossed to BDSC 458 for pan neuronal expression (See schematic Fig). Flies eclosed from the cross were collected in fresh food vials and kept under standard conditions for 1-2 days for acclimatization. Flies were separated into two vials for Mock buffer and F2,6BP treatment. One BD syringe (1 mL) was filled with Mock buffer, and the other was filled with F2,6BP and drop is administered per day to respective cohorts at the same time of day for 21 days.

### Climbing or Negative Geotaxis assay

The climbing assay was performed as described with minor modifications (Chakraborty *et al*., 2024). Briefly, experimental flies were anesthetized on ice. A group of 10 male flies per vial were transferred to a 25 mL sterile glass measuring cylinder. The measuring cylinder was divided into 6 compartments equally, the lowest compartment was labelled with 1 and the highest compartment was labelled with 6. The measuring cylinder with flies were placed against a white background for better video recording. The cylinder was tapped gently 3 times to send the flies to the bottom of the cylinder. The climbing time was recorded for 20s. Five trials were performed for each cohort. Two time-points, 8s and 16s were used to calculate the mean climbing score, which is total score divide by the total number of flies.

### Statistical analysis

All values shown on graphs represent mean ± standard deviation (SD). Statistical difference of means between data groups was determined using one-way ANOVA followed by Tukey’s range test. A p-value of p<0.05 was considered statistically significant.

## Acknowledgements

This work was supported by grants NIH/AG072487 and NIH/AI163327 to GG, and a grant from Cure Alzheimer’s Foundation to GG and JS; R56NS073976 to TH, funding and resources provided by the Moody Brain Health Institute at UTMB; MICIN/AEI/10.13039/501100011033 (PID2023-148933OB-I00 and RED2024-154053-T), CIBERCV (CB16/11/00222) from the Instituto de Salud Carlos III (co-financed by the European Development Regional Fund “A Way to Achieve Europe”, by the “European Union”; Intramural-CSIC 202320E201, and Comunidad de Madrid, Programa Biociencias (S2022-BMD-7223) to LB. NIH/NIA 4 R37 AG072502, and 5 P01 AG051449, the Bright Focus Foundation, and the Salk Stem Cell Core to FHG. Postmortem brain tissues and data from patients included in this study were provided three sources: UCSD Shiley Marcos Alzheimer’s Disease Research Center (ADRC), Mayo Clinic Brain Bank, Jacksonville, Florida and Banco de Tejidos CIEN (PT17/0015/0014), which has been integrated into the Spanish National Biobanks Network. We want to particularly acknowledge Dr Doug Galasko of ADRC for AD samples, Dr. Dennis Dickson for CBD and PSP samples. Analysis of published RNA-seq data used the AD Knowledge Portal (adknowledgeportal.org). These data were generated from postmortem brain tissue collected through the Mount Sinai VA Medical Center Brain Bank and were provided by Dr. Eric Schadt from the Icahn School of Medicine at Mount Sinai. We thank Dr. S. Jati for generating some data while he was working in the Ghosh laboratory as a postdoctoral fellow. We want to acknowledge UCSD Nikkon Microscopy Facility (Grant P30 NS047101) for support.

