## Supplementary Figures for "A Toxic Tau-PFKFB3 Circuit Reduces F2,6BP Levels and Drives Neurodegeneration"

**Supplementary Information**

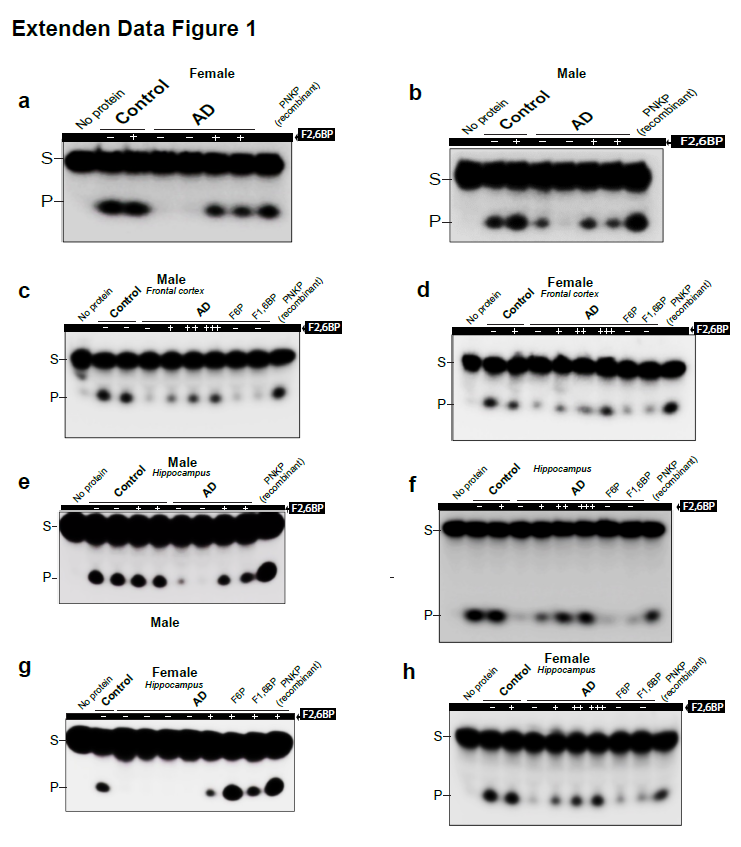

**Figure S1. Levels of F2,6BP, 3’-phosphatase activity of PNKP in AD and tauopathy patients.**

**a&b.** Representative autoradiograms illustrating 3’-phosphatase activity in nuclear extracts of frontal cortices from control and AD patients (G, female; H, male) with and without the supplementation of F2,6BP (50 𝜇M).

c. DNA 3’-phosphatase activity assay of PNKP in the nuclear extracts of frontal cortices from

representative control and AD patient samples (male) on a 32P-labeled 3’-phosphate containing oligo substrates mimicking SSBs. 2 ng pure recombinant PNKP was used as a

positive control. S= substrate and P= released phosphate. 3’-phosphatase activity of PNKP in

cortices of control and AD patient samples.

d. 3’-phosphatase activity of PNKP in hippocampi of male control and AD patient samples.

e&g. 3’-phosphatase activity of PNKP in hippocampi of female control and AD patient samples.

f&h. Dose dependent effect of F2,6BP on 3’-phosphatase activity in hippocampal nuclear from male (f) and female (h) control and AD patient samples. NP (no protein) and rPNKP (recombinant PNKP) are negative and positive control, respectively. F6P and F1,6BP showed no effect on the phosphatase activity in the nuclear extract.

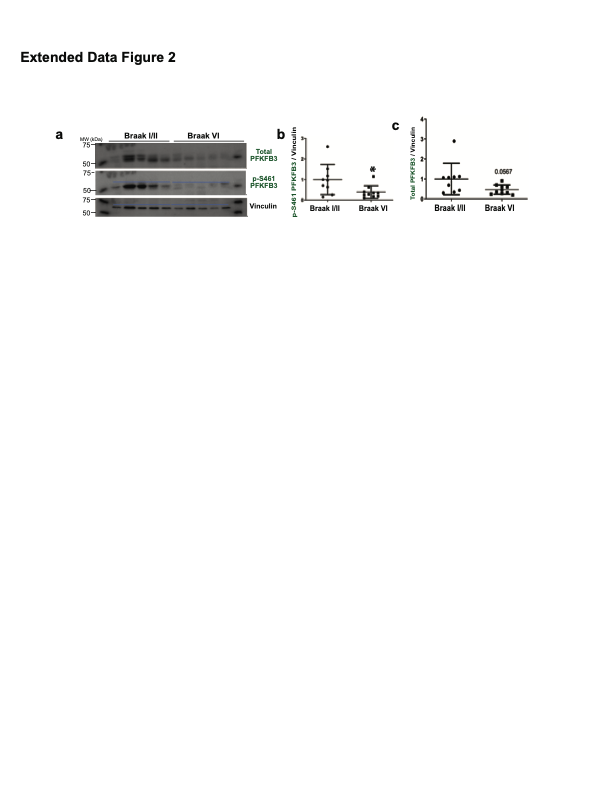

**Figure S2. Levels of F2,6BP, PFKFB3 and phosphor(p461)-PFKFB3 in patients.**

a.Western blot of showing PFKFB3 levels. Total PFKFB3, phospho-PFKFB3 and Vinculin levels using highly specific antibodies.

b&c. Normalized (with vinculin) levels of phosph-PFKFB3 (l) total PFKFB3 (j) and between AD and control samples.

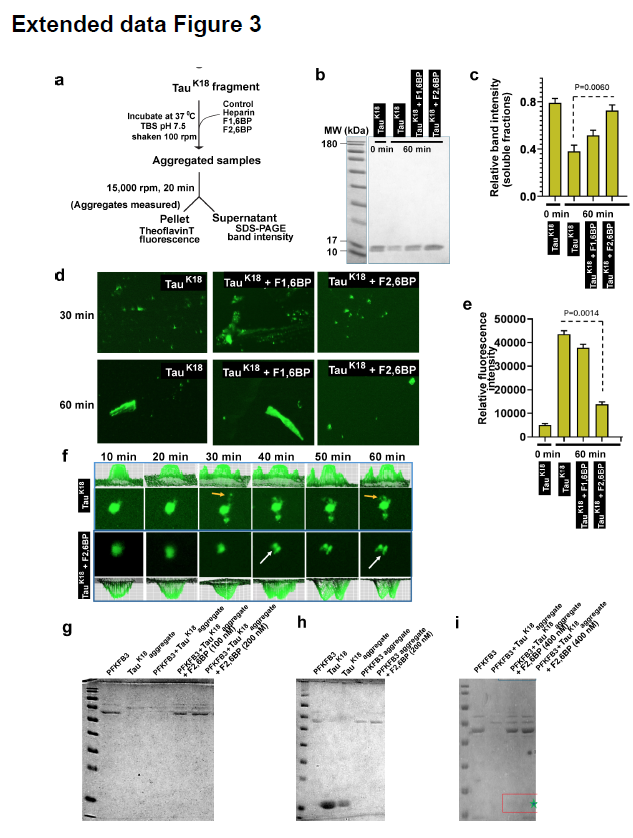

**Figure S3. F2,6BP prevents aggregation of K18, PFKFB3 and Myc-K18:PFKFB3 complex.**

**a.** Reaction scheme the experimental set up to monitor the effect of F2,6BP on Tau under different conditions (for experiments shown in Extended Data Figures 4B-E).

**b.** Representative SDS-PAGE showing levels of K18 after incubat of TauK18 for 0 min and with F1,6BP, F2,6BP or mock for 60 min. (n=3)

**c.** Quantitation mean of band intensities of three independent experiments ± SEM, shown in panel B.

**d.** Pellets from reactions in panel C were resuspended in buffer visualization by ThT fluorescence under an EVOS fluorescence microscope.

**e**. Quantitation of relative fluorescence intensities mean from three independent experiments ±SEM as shown in panel D.

**f**. F2,6BP disassembles Tau aggregates. Tracking of fluorescently labeled aggregates shows that F2,6BP disassembles the aggregate (white arrow). Continuous live imaging of Tau aggregates was performed in the presence and absence of F2,6BP. Fluorescently labelled TauK18 was monitored over time under in vitro conditions (TBS, pH 7.5, 37 °C) to assess aggregate formation and dynamics. The top row shows Tau aggregation in the absence of F2,6BP, where fluorescence intensity progressively increases and punctate aggregates form and grow over time (red arrows). The bottom row depicts Tau in the presence of F2,6BP, showing altered aggregation dynamics with smaller or dispersed aggregates and reduced fluorescence clustering. Each panel includes corresponding 3D intensity profiles (top and bottom of each row) illustrating the distribution and relative intensity of aggregates.

**g-i**. SDS-PAGE analysis of PFKFB3 and TauK18 aggregation independently, their combination and their prevention by F2.6BP.

**g.** Lane 1. MW standard. Lane 2. Fresh PFKFB3 (~60 kDa), Lane 3. Aggregated TauK18 (TauK18 aggregate) (not visible), Lane 4. TauK18 aggregate + fresh PFKFB3, 1 hr incubation before loading (PFKFB3 disappears). Lane 5. TauK18 aggregate + fresh PFKFB3 +100 nM F2,6BP, 1 hr incubation (PFKFB3 reappears) Lane 6. TauK18 aggregate+fresh PFKFB3 +200 nM F2,6BP 1 hr incubation (more PFKFB3 reappears).

**h.** Lane 1. MW standard. Lane 2. Fresh PFKFB3, Lane 3. Fresh Tau, Lane 4. Fresh TauK18 incubated for 1 hr before loading (TauK18 aggregate), Lane 5. PFKFB3 incubated for 1 hr before loading (PFKFB3 partially disappears). Lane 6. PFKFB3 + 200 nM F2,6BP incubated for 1 hr before loading (PFKFB3 reappears).

**i.** Lane 1. MW standard, Lane 2. Fresh PFKFB3, Lane 3. PFKFB3 + TauK18 aggregate incubated 1 hr, Lane 4. PFKFB3 + TauK18 aggregate + 200 nM F2,6BP incubated 1 hr , Lane 5. PFKFB3 + TauK18aggregate + 400 nM F2,6BP incubated 1 hr.

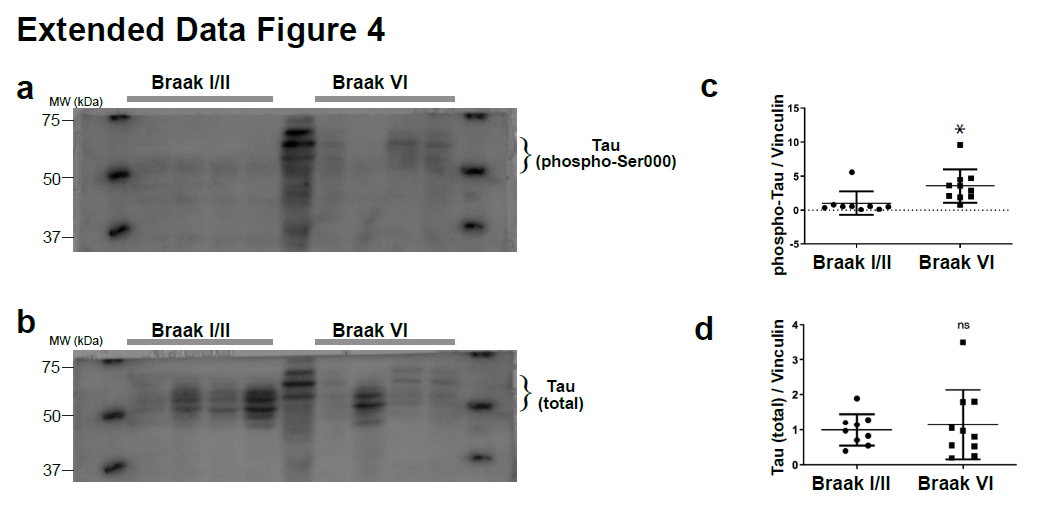

**Figure S4. Relative levels of phospho- and total tau in AD and control samples.**

a. Representative western blot showing levels of phospho-Tau in AD and control samples.

b. Representative western blot showing levels of total Tau in AD and control samples

c. Normalized (with Vanculin) levels of phosph-Tau in patient (n= 10) versus control (n=9) samples. “*” denotes significance levels <0.05.

d. Normalized (with Vanculin) levels of total Tau in patient (n=10) versus control (n=9) samples.

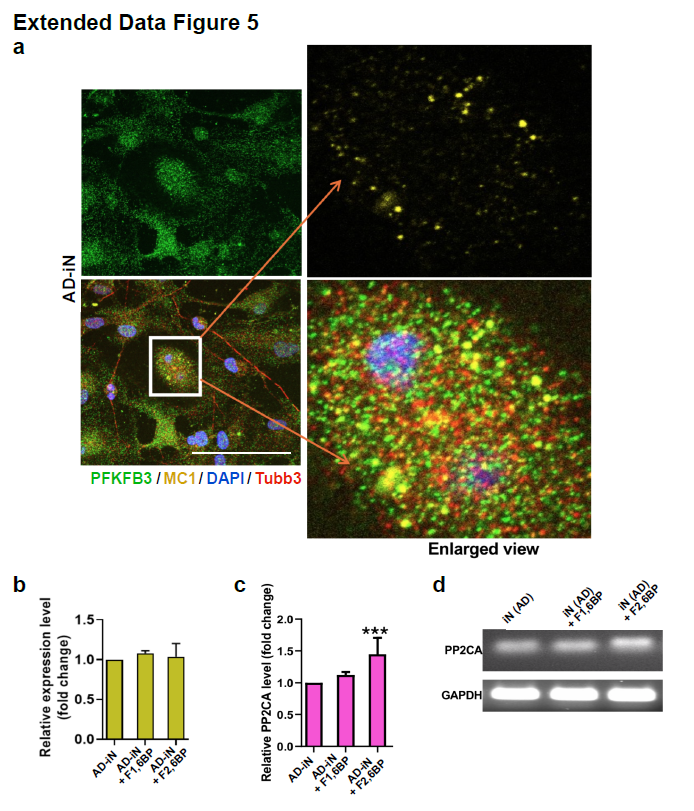

**Figure S5. Tau-PFKFB3 colocalization and transcriptional effect upon F2,6BP treatment**

**a.** Close-up view of confocal images of AD-iNs stained with indicated antibodies to confirms the Tau- PFKFB3 colocalization.

**b.** RT-qPCR analysis of PFKFB3 mRNA levels in AD-iNs untreated and treated with indicated

molecules **c.** RT-qPCR analysis of pp2ca mRNA levels in AD-iNs untreated and treated with F2,6BP.

**d.** The same qPCR reactions were run and visualize on an agarose gel.

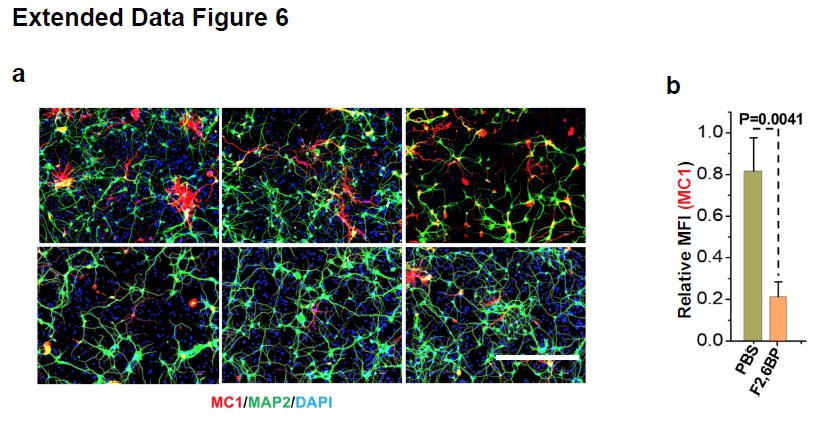

**Figure S6. The effect of F2,6BP on AD-Primary neurons.**

**a. T**he effect of F2,6BP on primary neurons and staining with MC1 and MAP2 antibodies and imaging after F2,6BP treatment, PBS (control, top) treatment (Bottom).

**b.** Quantification of MC1 and MAP2 staining. Scale bar=50µm.
