## Supplementary Information for "A Toxic Tau-PFKFB3 Circuit Reduces F2,6BP Levels and Drives Neurodegeneration"

**Supplementary Information *(SI APPENDIX)***

**Supplementary Table 1**

| H-TUBB3-LA-F | TUBB (human) | TGCTTCTCATGCTTGCTACCAC | LA-qPCR |
| --- | --- | --- | --- |
| H-TUBB3-LA-R | TUBB (human) | TCTGTCCCTGTAGGAGGATGT | LA-qPCR |
| H-NeuroD1-LA-F | NeuroD1 (human) | CCGCGCTTAGCATCACTAAC | LA-qPCR |
| H-NeuroD1-LA-R | NeuroD1 (human) | TGGCACTGGTTCTGTGGTATT | LA-qPCR |
| H-ENO2-LA-F | Enolase (human) | ACGTGTGCTGCAAGCAATTT | LA-qPCR |
| H-ENO2-LA-R | Enolase (human) | CCTGAAACTCCCCTGACACC | LA-qPCR |
| H-MyH2-LA-F | MyH2 (human) | AAAGCCTGCCAAGCCCTAAA | LA-qPCR |
| H-MyH2-LA-R | MyH2 (human) | TGGTCAGCATGGCAAGTGAA | LA-qPCR |
| H-MyH4-LA-F | MyH4 (human) | CAGGAGTGGTCCCTAAAGGC | LA-qPCR |
| H-MyH4-LA-R | MyH4 (human) | GTAAAAACACGGTCCCTGCC | LA-qPCR |
| H-MyH6-LA-F | MyH6 (human) | ATTGACTCGGTGCCCTTTCT | LA-qPCR |
| H-MyH6-LA-R | MyH6 (human) | TGCCACCCAACATGGGTATT | LA-qPCR |
| H-TUBB Primer mix | TUBB (human) | RealTimePrimers.com | SA-PCR/qRT-PCR |
| H-Enolase Primer mix | Enolase (human) | RealTimePrimers.com | SA-PCR/qRT-PCR |
| H-MyH2 Primer mix | MyH2 (human) | RealTimePrimers.com | SA-PCR |
| H-MyH4 Primer mix | MyH4 (human) | RealTimePrimers.com | SA-PCR |
| H-MyH6 Primer mix | MyH6 (human) | RealTimePrimers.com | SA-PCR |
| **Oligo sets for amplification of mitochondrial fragment from *Drosophila* genome:** | | | |
| LA 1: 5’-TGTGAATAATAGCCCCAGCACA-3’ | | |  |
| LA 2: 5’-GCTGGAATGAATGGTTGGACG-3’ | | |  |
| SA 1: 5’-ACACCTGCCCATATTCAACCA-3’ | | |  |
| SA 2: 5’-ACTGGTCGAGCTCCAATTCA-3’ | |  |  |
| **qPCR primers** |  |  |  |
| ppp2ca_F | CAGTGGATCGAGCAGCTGAA | |  |
| ppp2ca_R | TTTGTCAGGATTTCTTTAGCCTTC | |  |
| pfkfb3_F | CAGCTGCCTGGACAAAACAT | |  |
| pfkfb3_R | CGTCTGCCTCAGTGTTTCCT | |  |
| gapdh-rtpcr-f | GTCTCCTCTGACTTCAACAGCG | |  |
| gapdh-rtpcr-r | ACCACCCTGTTGCTGTAGCCAA | |  |

**Patient Information**

| PATHID | PATHDX1 | BRAAK | Death | Age | Sex | APOE |
| --- | --- | --- | --- | --- | --- | --- |
| 5919 | Normal | 1 | 12/13/20 | 75 | F | 23 |
| 5856 | Alzheimers changes | 2 | 10/2/18 | 83 | M | 33 |
| 5844 | Normal | 2 | 3/6/18 | 96 | F | 33 |
| 5783 | Normal | 2 | 10/19/16 | 84 | M | 23 |
| 5747 | Alzheimers changes | 2 | 12/1/15 | 88 | M | 34 |
| 5709 | Normal | 2 | 11/17/14 | 94 | M | 33 |
| 5699 | Alzheimers changes | 2 | 8/28/14 | 92 | F | 33 |
| 5687 | Normal | 1 | 5/30/14 | 84 | M | 33 |
| 5662 | Alzheimers changes | 2 | 11/12/13 | 93 | F | 33 |
| 5655 | Alzheimers changes | 2 | 9/23/13 | 82 | F | 34 |
| 5567 | Alzheimers changes | 1 | 2/1/12 | 86 | M | -4 |
| 5546 | Alzheimers changes | 1 | 7/7/11 | 91 | F | 33 |
| 5515 | Alzheimers changes | 1 | 9/4/10 | 73 | F | -4 |
| 5517 | Alzheimers changes | 1 | 10/9/10 | 86 | M | -4 |
| 5529 | Alzheimers changes | 1 | 2/15/11 | 81 | M | 33 |
| 5447 | Alzheimers changes | 1 | 4/9/09 | 91 | F | -4 |
| 5398 | Alzheimers changes | 0 | 5/2/08 | 89 | F | -4 |
| 5687 | Normal | 1 | 5/30/14 | 84 | M | -4 |
| 5971 | Alzheimers disease | 6 | 1/7/23 | 74 | F | 44 |
| 5970 | Alzheimers disease | 6 | 12/24/22 | 66 | M | 34 |
| 5969 | Alzheimers disease | 6 | 12/16/22 |  | M |  |
| 5966 | Alzheimers disease | 6 | 12/3/22 |  | F |  |
| 5965 | Alzheimers disease | 6 | 9/9/22 | 75 | F | 34 |
| 5964 | Alzheimers disease | 6 | 9/6/22 | 91 | M | 34 |
| 5959 | Alzheimers disease | 6 | 7/21/22 | 70 | M | 34 |
| 5958 | Alzheimers disease | 6 | 6/12/22 | 86 | F | 34 |
| 5953 | Alzheimers disease | 6 | 2/25/22 | 58 | M | 34 |
| 5952 | Alzheimers disease | 6 | 2/2/22 | 79 | F | 34 |
| 5741 | Alzheimers disease | 6 | 10/13/15 | 90 | F | -4 |
| 5740 | Alzheimers disease | 6 | 10/13/15 | 88 | F | -4 |
| 5738 | Alzheimers disease | 6 | 10/3/15 | 85 | F | -4 |
| 5736 | Alzheimers disease | 6 | 8/1/15 | 65 | F | -4 |
| 5750 | Alzheimers disease | 6 | 1/18/16 | 85 | F | -4 |
| 5748 | Alzheimers disease | 6 | 12/7/15 | 68 | F | -4 |
| 5759 | Alzheimers disease | 6 | 2/25/16 | 87 | F | -4 |
| 5755 | Alzheimers disease | 6 | 2/9/16 | 83 | F | -4 |
| 5753 | Alzheimers disease | 6 | 1/31/16 | 67 | F | -4 |
| 5774 | Alzheimers disease | 6 | 7/13/16 | 87 | F | -4 |
| 5770 | Alzheimers disease | 6 | 5/24/16 | 84 | M | -4 |
| 5768 | Alzheimers disease | 6 | 5/5/16 | 69 | M | -4 |
| 5767 | Alzheimers disease | 6 | 4/15/16 | 92 | M | -4 |
| 5788 | Alzheimers disease | 6 | 11/19/16 | 92 | M | -4 |
| 5779 | Alzheimers disease | 6 | 9/10/16 | 73 | M | -4 |
| 5799 | Alzheimers disease | 6 | 2/2/17 | 82 | F | -4 |
| 5797 | Alzheimers disease | 6 | 1/19/17 | 81 | M | -4 |
| 5796 | Alzheimers disease | 6 | 1/9/17 | 96 | M | -4 |
| 5795 | Alzheimers disease | 6 | 1/6/17 | 81 | M | -4 |
| 5789 | Alzheimers disease | 6 | 11/26/16 | 89 | M | -4 |
